# Universal protein misfolding intermediates can bypass the proteostasis network and remain soluble and less functional

**DOI:** 10.1101/2021.08.18.456613

**Authors:** Daniel Nissley, Yang Jiang, Fabio Trovato, Ian Sitarik, Karthik Narayan, Philip To, Yingzi Xia, Stephen D. Fried, Edward P. O’Brien

## Abstract

Misfolded protein conformations with decreased functionality can bypass the proteostasis machinery and remain soluble *in vivo*. This is an unexpected phenomenon as several cellular quality control mechanisms have evolved to rid cells of misfolded proteins. Three questions, then, are: how is it structurally possible for long-lived, soluble, misfolded proteins to bypass the proteostasis machinery and processes? How widespread are these soluble, misfolded states across the proteome? And how long do they persist for? Here, we address these questions using coarse-grain molecular dynamics simulations of the synthesis, termination, and post-translational dynamics of a representative set of cytosolic *E. coli* proteins. We predict that half of all proteins exhibit subpopulations of misfolded conformations that are likely to bypass molecular chaperones, avoid aggregation, and not be rapidly degraded. These misfolded states may persist for months or longer for some proteins. Structurally characterizing these misfolded states, we observe they have a large amount of native structure, but also contain localized misfolded regions from non-native changes in entanglement, in which a protein segment threads through a loop formed by another portion of the protein that is not found in the native state. The surface properties of these misfolded states are native like, suggesting they may bypass the proteostasis machinery and its regulatory processes to remain soluble, while their entanglements make these states long-lived kinetic traps, as disentanglement requires unfolding of already folded portions of the protein. In terms of function, we predict that one-third of proteins have subpopulations that misfold into less-functional states that have structurally perturbed functional sites yet remain soluble. Data from limited-proteolysis mass spectrometry experiments, which interrogate the misfolded conformations populated by proteins upon unfolding and refolding, are consistent with the structural changes seen in the entangled states of glycerol-3-phosphate dehydrogenase upon misfolding. These results provide an explanation for how proteins can misfold into soluble conformations with reduced functionality that can bypass cellular quality controls, and indicate, unexpectedly, this may be a wide-spread phenomenon in proteomes. Such entanglements are observed in many native structures, suggesting the non-native entanglements we observe are plausible. More broadly, these near-native entangled structures suggest a hypothesis for how synonymous mutations can modulate downstream protein structure and function, with these mutations partitioning nascent proteins between these kinetically trapped states.

## INTRODUCTION

How soluble, misfolded protein populations with reduced functionality^1–3^ bypass cellular quality control mechanisms^4^ for long time periods is poorly understood. Further, how common this phenomenon is across organismal proteomes has not been assessed. These are important gaps in our knowledge to fill as the answers will offer a more complete picture of protein structure^5^ and function in cells, may lead to refinement of the protein homeostasis model^6^ of proteome maintenance, are likely to be relevant to how synonymous mutations have long-term impacts on protein structure and function^7^, and could reveal long-term misfolding on a scale greater than previously thought.

Protein homeostasis (“proteostasis”) refers to the maintenance of proteins at their correct concentrations and in their correct conformational states through the action of a cohort of chaperones, degradation machineries, and protein quality control pathways^6,8^. It is typically posited that under normal (*i.e*., not stressed) cellular growth conditions globular proteins *in vivo* attain one of three states: folded/functional, misfolded/aggregated, or degraded. Various molecules work together to maintain proteostasis by catalyzing the interconversion of proteins between these states^9–11^. For example, some chaperones in *E. coli*, such as GroEL/GroES^9^ and DnaK^10^, can promote the folding of misfolded or unfolded proteins. Others, such as the set of enzymes associated with the ubiquitin-proteasome system in eukaryotes, covalently tag misfolded proteins for degradation^12^. Yet others, such as *E. coli*’s ClpXP, have the potential to break apart aggregates, allowing released monomeric proteins to be degraded^13^. Many caveats and nuances exist in the proteostasis model. For example, some insoluble protein aggregates, such as carboxysomes, are biologically beneficial by spatially concentrating protein function^14^. Some non-native protein oligomers are soluble^15^, and recently discovered biomolecular condensates^16^ represent a form of phase separation in which proteins within a condensate remain soluble but preferentially interact with each other over other cellular components. Additionally, soluble proteins are not always functional because some require co- or post-translational modifications^17,18^.

The timescales involved with many proteostasis processes are often quite short. Co-translationally acting chaperones bind ribosome nascent chain complexes on time scales of tens of ms^[19]^, the ubiquitin-degradation machinery tags up to 30% of eukaryotic nascent chains for immediate degradation after synthesis^20,21^, and post-translationally acting chaperones can generally refold misfolded proteins in seconds or minutes^22^. Indeed, the FoldEco kinetic model of *E. coli* proteostasis indicates that conversions between various states within the network occur with rate constants typically on the order of seconds^23^. Thus, according to the proteostasis model, misfolded proteins should either be converted in a matter of seconds or minutes into their folded state, be degraded, or form aggregates provided cells are not stressed and the proteostasis machinery is not overwhelmed^24^.

Synonymous mutations change the triplet of nucleotides between degenerate mRNA codons encoding the same amino acid, leading to an altered mRNA sequence that encodes the same protein primary structure. Such mutations can alter the translation-elongation rate of ribosomes and have been found to alter the structure and function of proteins for long timescales^7^. For example, translation of a synonymous variant of the *frq* gene in the fungus *Neurospora* resulted in the synthesis of FRQ protein with altered conformations that bound 50% less to a partner protein, resulting in a significantly altered circadian rhythm that persisted for multiple days^2^.

Many other proteins have been reported to exhibit altered structure or function upon the introduction of synonymous mutations^25–27^. The fact that these functional changes occur in the soluble fraction of the proteome indicates it is not insoluble aggregation driving this phenomenon. And importantly, such observations are inconsistent with aspects of the proteostasis model, which predicts that any protein with a misfolded (and less functional) structure should either refold, aggregate, or be degraded on faster timescales.

One hypothesis that could resolve this discrepancy is that proteins can populate an additional state. In this state, proteins are kinetically trapped over long timescales in misfolded conformations with reduced functionality, but they do not have a propensity to aggregate or interact with proteostasis machinery in excess of that of folded proteins. If correct, this hypothesis raises a number of questions, including: (*i*) what type of structures are adopted in this state? (*ii*) how do those conformations simultaneously avoid folding, aggregation, and degradation in excess of that observed for the native ensemble? (*iii*) how long do they persist? And (*iv*) what fraction of the proteome exhibits this behavior?

Answering these questions requires a computational method that can access the second to minute timescale of protein synthesis and maturation while providing sufficient structural resolution to identify misfolded conformations and their properties. We use a topology-based coarse-grain model that represents proteins with one interaction site per residue placed at the coordinates of the Cα atom. This model folds proteins 4-million times faster, on average, than in real systems^28^, and was previously used to accurately reproduce the co-translational folding time course of HemK N-terminal domain^29^. Coarse-grain simulations of a zinc-finger protein folding in the ribosome exit tunnel were also found to agree with experimental cryo-EM structures^30^, indicating this method can reproduce realistic scenarios of folding on the ribosome. Additionally, excellent agreement has been found between such topology-based models and experimental assays monitoring force generation due to the folding of titin I27 domain on and off the ribosome^31^. These examples highlight the utility of such coarse-grain models to protein misfolding on and off the ribosome.

Here, we use such coarse-grain methods to simulate protein synthesis, co-translational and post-translational folding, and estimate the fraction of molecules that fold, misfold, interact with chaperones, aggregate, are degraded, or attain a functional conformation. After first confirming that our model can reproduce post-translational misfolding in Luciferase, we simulate a representative subset of the cytosolic *E. coli* proteome, finding that a substantial proportion of newly synthesized proteins can adopt misfolded conformations that are near-native in structure and thus likely to interact with co- and post-translational chaperones in a manner similar to that of their native states. These misfolded conformations expose a similar amount of aggregation-prone surface area as the native ensemble, and therefore do not have an increased propensity to aggregate. For some proteins, misfolding is localized near their functional sites, indicating their functionality is reduced. We estimate that many of these near-native misfolded states are kinetically trapped, exhibiting lifetimes on the order of days to months. Our simulations predict that there is a universal structural feature of these proteome-wide, soluble misfolded states.

## RESULTS

### A coarse-grain model reproduces experimentally observed misfolding of Firefly Luciferase

The model we use for protein synthesis, folding, and function has been shown to accurately predict experimentally measured changes in enzyme specific activities^32^, indicating it reasonably describes protein structure-function relationships. As an additional test, here we examine if the model is able to identify if a protein will exhibit misfolded subpopulations. Firefly Luciferase, a 550-residue protein with four domains, folds co-translationally^33^. Specific activity experiments^25^ have found that some soluble, nascent Luciferase molecules misfold when translation speed is increased. Even when synthesized from its wild-type mRNA in *E. coli* some Luciferase molecules still fail to fold correctly^25^. We therefore selected Luciferase as a test system, judging that if it partitions into long-lived misfolded states in our simulations when translated from its wild-type mRNA that our model is able to capture realistic scenarios of misfolding.

We simulated Luciferase’s synthesis, ejection from the ribosome exit tunnel, and post-translational dynamics using a coarse-grain representation of the protein and ribosome (Figure 1a-c, Table S1, Methods, and Eq. 1 and 2). Fifty statistically independent trajectories were run. To characterize Luciferase’s native conformational ensemble we also simulated ten trajectories initiated from Luciferase’s crystal structure, which we refer to as “native-state simulations”. To assess whether or not a given Luciferase trajectory is misfolded we utilize the time-dependent mode of the fraction of native contacts (*Q*_mode_) and the probability that a non-native entanglement (*P*(*G_k_*)) has formed. We categorize a trajectory as misfolded if it has (*i*) a mean *Q*_mode_, over the final 100 ns of the post-translational phase of the simulation, that is less than the average from the native-state simulations, or (*ii*) a mean *P*(*G_k_*) for the different possible changes in non-covalent lasso threading denoted *k* = {0,1,2,3,4} of 0.1 or greater over the final 100 ns of the trajectory, or (*iii*) both (*i*) and (*ii*) occur (see Methods). Conditions (*i*) and (*ii*) correspond to perturbations of structure relative to the native state as defined by the fraction of native contacts and entanglement, respectively. Based on this definition, 46% (95% Confidence Interval [32%, 60%], calculated from bootstrapping 10^6^ times) of nascent Luciferase molecules misfold (Figure 1d-e, Methods). We observe that when Luciferase misfolds, it misfolds 100% of the time in the second domain (based on 〈*Q*_mode_〉), which is composed of residues 13-52 and 212-355. These misfolded Luciferase structures are near-native, with a 5.5% decrease in the overall fraction of native contacts (〈*Q*_overall_〉 = 0-86, computed over the final 100 ns of misfolded trajectories) compared to the native ensemble (〈*Q*_overall_〉 = 0-91). Thus, a large proportion of nascent Luciferase misfolds into near-native conformations that typically involve misfolding of the second domain.

**Figure 1.**
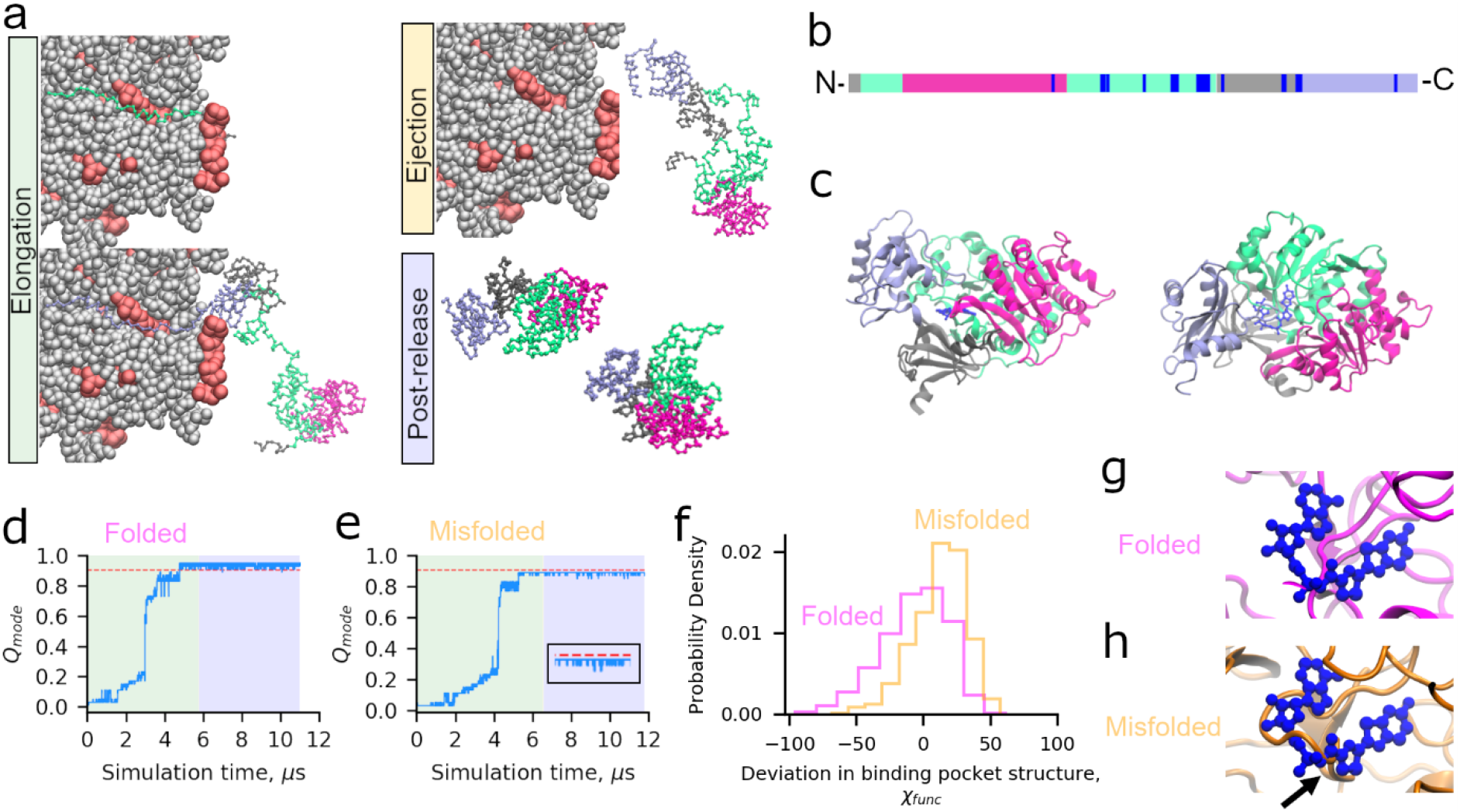
Luciferase exhibits subpopulations that misfold into soluble but less-functional conformations. (a) Simulations of translation elongation and ejection of nascent Luciferase were performed with a coarse-grain ribosome representation (ribosomal proteins and RNA are displayed in red and grey respectively). Domains 1, 2, 3, and 4 of Luciferase are displayed in silver, light green, magenta, and light purple, respectively. After ejection, the ribosome is removed and post-release dynamics is simulated for 30 CPU days per trajectory. (b) Primary structure diagram of Luciferase colored as described in (a); positions involved in the catalytic function of Luciferase as described in Methods are colored blue. (c) Cartoon diagram of Luciferase native state colored as described in (a) with the 5′-O-[N-(dehydroluciferyl)-sulfamoyl]-adenosine ligand colored dark blue. (d) *Q*_mode_ (see Methods) versus time for Domain 2 of a trajectory of Luciferase that folded correctly. Portions of the plot colored green, yellow, and blue correspond to the synthesis, ejection, and post-translation phases of the simulation. Note that the relatively short duration of ejection for this protein renders that section of the plot invisible at this resolution. The red line corresponds to 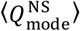 minus three standard deviations and represents the threshold for defining this domain as folded (see Methods). (e) Same as (d) but for a trajectory that misfolds. Inset shows the final microsecond of the *Q*_mode_ time series. (f) Distributions of *χ*_func_ (Eq. 13) over the final 100 ns of the folded (magenta) and misfolded (orange) trajectories displayed in panels (d) and (e). The misfolded and folded distributions are different based on the Kolmogorov-Smirnov test with test statistic 0.33 and *p*-value of 1 x 10^−66^. The misfolded distribution shows greater structural distortion (i.e., values of *χ*_func_> 0) of the binding pocket. (g) Backmapped all-atom structure from the final frame of the folded simulation shown in (d) aligned based on the residues implicated in function to the native state. (h) Same as (g) except for the final structure from the misfolded trajectory in (e), showing a strand misfolding in the ligand binding pocket (indicated by black arrow). Steric conflict between where the substrate binds and the surrounding binding pocket of the misfolded structures indicates this misfolded state will have reduced function.

The motivating experiments on Luciferase were carried out in the presence of the endogenous *E. coli* proteostasis machinery^25^. To predict whether the Luciferase misfolded states produced by our model are likely to display reduced specific activity *in vivo* we therefore need to determine four things. They must (*i*) evade chaperones to remain misfolded, (*ii*) not aggregate, (*iii*) not get degraded, and (*iv*) the residues involved in function must be structurally perturbed. The chaperone trigger factor (TF) binds nascent proteins co-translationally, DnaK interacts both co- and post-translationally, while GroEL/GroES is primarily a post-translational chaperone. Interactions with TF^34^ or GroEL/GroES^9^ are thought to occur by the non-specific recognition of exposed hydrophobic patches on client proteins co- and post-translationally, respectively. DnaK, however, is hypothesized to interact with specific binding sites within a protein’s sequence^35^.

To estimate whether misfolded Luciferase is likely to interact with TF we computed the average relative difference between the hydrophobic solvent accessible surface area (SASA) of each misfolded trajectory to the folded population (denoted 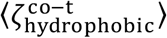, Eq. 9 and Methods) during synthesis. The value of 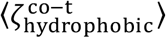 is ≤10% for 16 of 23 misfolded trajectories, meaning that they display less than a 10% increase in hydrophobic SASA during synthesis relative to the folded population of trajectories. This indicates that a majority of misfolded Luciferase molecules will not interact with TF much more than a properly folded Luciferase molecule (see Methods). Though TF accelerates protein folding under force^36^, under normal conditions it is also thought to act as a holdase^34^. Thus, we conclude that our co-translational Luciferase misfolded states can misfold into conformations that do not interact with TF in a manner that accelerates folding, allowing these misfolded states to persist post-translationally.

Next, to determine whether misfolded conformations of Luciferase are likely to interact with GroEL/GroES post-translationally, we computed the average relative difference between the hydrophobic SASA of each misfolded conformation in the final 100 ns and its value in the native-state simulations (〈*ζ*_hydrophobic_〉, Eq. 10, Methods). The value of 〈*ζ*_hydrophobic_〉 for Luciferase is ≤10% for 9 of 23 misfolded trajectories, indicating that these misfolded states expose only a small excess of hydrophobic SASA relative to the native ensemble, and are therefore not likely to be engaged by GroEL/GroES.

Finally, to estimate whether misfolded Luciferase structures are more likely to interact with DnaK than the native state, we computed 〈*ζ*_DnaK_〉 (Eq. 11), the average relative difference in SASA of residues predicted to be DnaK binding sites by the Limbo algorithm^35^ in the final 100 ns for all misfolded trajectories. We find that 22 out of 23 misfolded trajectories have 〈*ζ*_DnaK_〉 ≤ 10%, indicating that DnaK is unlikely to preferentially bind to these misfolded states any more than it is to the native state. Thus, some of Luciferase’s misfolded states are unlikely to interact with TF, GroEL/GroES, or DnaK, and thus bypass the *E. coli* chaperone network (Figure S1a).

The next key question is whether or not these misfolded Luciferase structures, having bypassed chaperone quality controls, are likely to remain soluble or to aggregate or be degraded. In the original experiments by Barral and co-workers, centrifugation was used to remove aggregates from the soluble fraction. To estimate whether the misfolded Luciferase structures from our simulations will aggregate we used the AMYLPRED2^37^ webserver to identify residues in the Luciferase amino acid sequence that lead to aggregation when exposed to solvent. We then computed 〈*ζ*_agg_〉 (Eq. 12), the average relative difference in SASA between these aggregation-prone residues in the final 100 ns of each misfolded trajectory in comparison to the same residues in the native-state simulations. We find that 12 of 23 misfolded trajectories have 〈*ζ*_agg_〉 ≤ 10%, indicating that these misfolded conformations display only a minor increase in aggregation propensity and are likely to remain soluble.

Finally, we considered the likelihood that misfolded Luciferase will be targeted for degradation. Degradation in *E. coli* is carried out primarily by proteases coupled to AAA+ ATPase motor proteins^38^, including ClpXP and Lon, that recognize and degrade misfolded or aggregated proteins. Misfolded protein structure contributes to degradation^39^, and therefore, like our GroEL/ES assessments, we use 〈*ζ*_hydrophobic_〉 to quantify how similar misfolded Luciferase conformations are to the native state. For 9 of 23 misfolded Luciferase trajectories 〈*ζ*_hydrophobic_〉 is ≤ 10%, indicating they are unlikely to be degraded more quickly than native Luciferase.

Having determined that some misfolded conformations of Luciferase can evade chaperones, aggregation, and degradation, the final question is whether their function is decreased relative to native Luciferase. To answer this question, we identified the residues that take part in Luciferase’s bioluminescence, defined as those resides within 4.5 Å of the 5’-O-[N-(dehydroluciferyl)-sulfamoyl]-adenosine ligand in PDB structure 4G36, in addition to all residues identified in the UniProt database^40^ to have a role in its catalytic mechanism. To quantify the difference in structure of residues involved in Luciferase’s catalytic mechanism, we compute the average relative difference between the structures sampled in the final 100 ns of each misfolded trajectory and native Luciferase in terms of the structural overlap function (〈*χ*_func_〉) over residues implicated in its function (see Eq. 13–15 and Figure 1b, c, f, g, and h). Positive values of *χ*_func_ indicate perturbed structure relative to the native state simulations. We find that 15 of 23 misfolded Luciferase trajectories have 〈*χ*_func_〉 ≥ 10%, indicating that they have significantly perturbed structure at functionally important sites relative to the native state (Figure 1f). For example, Figures 1g and 1h show the binding pocket at the final frames of folded and misfolded, soluble, but non-functional Luciferase trajectories. The binding pocket structure is perturbed such that it impinges on the substrate location. Since structure equals function, this result indicates that the efficiency of the enzymatic reaction carried out by misfolded Luciferase will be less efficient than in its native fold.

Cross-referencing the lists of misfolded trajectories that are likely to avoid chaperones, aggregation, degradation, and exhibit reduced function, we find that one trajectory displays all of these characteristics and likely remains soluble but less functional than native Luciferase (Figure S1b). Our simulation results are thus qualitatively consistent with the experimental observation that some nascent Luciferase molecules misfold when translated from its wild-type mRNA. While a misfolded state that is only populated by 2% of protein molecules is unlikely to strongly influence the cell, perturbations to Luciferase translation-elongation kinetics by synonymous mutations might increase this population beyond 2%. In general, these results indicate that our coarse-grain simulation protocol for nascent protein synthesis, ejection, and post-translational dynamics is able to recapitulate nascent protein misfolding.

### Simulating a representative subset of the *E. coli* cytosolic proteome

It is not computationally feasible to simulate all 2,600 cytosolic *E. coli* proteins. Therefore, to investigate the extent of nascent protein misfolding within the *E. coli* proteome we constructed models for a representative subset of 122 proteins. This set of proteins has the same distributions of protein length and structural class, and a similar ratio of multi- to single-domain proteins as the entire *E. coli* proteome^41^ (Table S2). The details of the parameterization of these models are described in Ref. 41. Each protein was synthesized on the same coarse-grain ribosome representation as Luciferase and their post-translational dynamics simulated for 30 CPU days per trajectory. Larger proteins take longer to simulate. Therefore, this fixed post-translational simulation run time resulted in trajectories of different durations due to different protein sizes. The simulation time in the post-translational phase therefore ranged between 2.7 and 154.1 *μ*s per trajectory. Because our coarse-grain model exhibits an approximately four-million fold acceleration of folding dynamics^28^, due to decreased solvent viscosity^42^ and a smoother free-energy landscape^43^, these post-translational simulation times correspond approximately to experimental times of 11.0 to 611 seconds, respectively. As with Luciferase, ten trajectories were also initiated from the crystal structure of each protein and simulated for 30 CPU days to serve as reference simulations representing the native-state structural ensemble.

### Two thirds of nascent *E. coli* proteins populate misfolded states

A fundamental question our simulation data set can address is how common nascent protein misfolding is across *E. coli*’s cytosolic proteome. As with Luciferase, we use the fraction of native contacts and entanglement as measures of misfolding. We find that 66% of proteins (80 out of 122) remain misfolded in at least one trajectory, 40% of proteins are misfolded in at least 20% of trajectories (49 out of 122), and 7% are misfolded in 100% of trajectories (9 out of 122). The proteins in these various categories are summarized in Table S4; Figure 2a displays a histogram of the probability of misfolding over the 122 different *E. coli* proteins simulated. In total, 27% of the simulation trajectories (1,631 out of 6,100) of the *E. coli* cytosolic proteome remain in misfolded conformations after 30 CPU days of post-translational dynamics.

**Figure 2.**
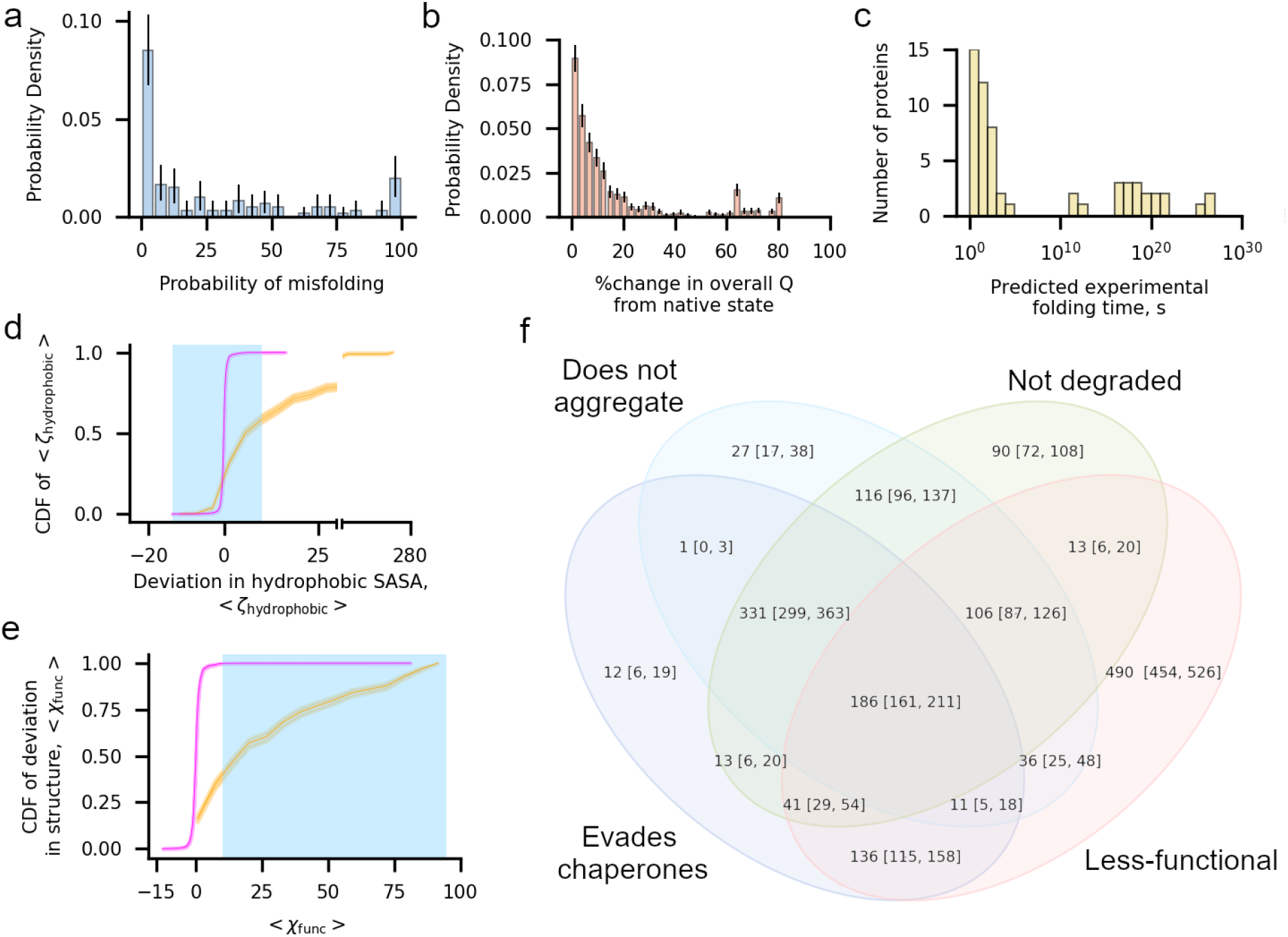
One in three proteins exhibit subpopulations that misfold into soluble but less-functional conformations that evade proteostasis machinery. (a) Histogram of the probability of misfolding being detected in the final 100 ns of the simulatio n, computed as the number of misfolded trajectories divided by 50, for each of the 122 proteins in the cytosolic *E. coli* proteome set. (b) Histogram of the percent change, computed as 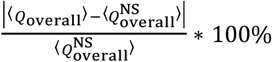, in fraction of native contacts within the final 100 ns of each of the 1,631 misfolded trajectories (〈*Q*_overall_〉) in the *E. coli* proteome data set relative to the average value from each protein’s native state simulations 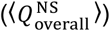. The majority of misfolded proteins are within 10% of the native value. (c) Histogram of extrapolated folding times for the slow-folding kinetic phase from survival probability curves for the 73 proteins in the cytosolic *E. coli* dataset with a reliable estimate (see Methods). (d) Cumulative distribution function (CDF) of 〈*ζ*_hydrophobic_〉 computed over the values of 〈*ζ*_hydrophobic_〉 (Eq. 10) for 1,631 misfolded (orange) and 4,469 folded (magenta) trajectories. The blue shaded region indicates the set of 〈*ζ*_hydrophobic_〉 values considered to have no significant increase in hydrophobic solvent-accessible surface area relative to the native-state ensemble. (e) Same as (d) but CDFs are computed over values of 〈*χ*_func_〉 for trajectories in the misfolded and folded populations. Blue shaded region indicates the set of values considered to result in perturbed function. (f) Venn diagram indicating the number of the 1,631 misfolded trajectories that evade chaperones (TF, DnaK, and GroEL/GroES), do not aggregate, are not degraded, and are non-functional. The 186 trajectories at the center of this diagram are misfolded states that are expected to evade the proteostasis machinery, remaining soluble but non-functional. All error bars are 95% confidence intervals computed from bootstrapping 10^6^ times; the height of the CDF plots in (d) and (e) indicates the 95% confidence intervals.

### Many misfolded states are similar to the native state

Misfolded conformations that are very different from the native state will likely interact with the proteostasis machinery. To characterize the closeness of misfolded states to the native state across our set of misfolded conformations we calculated the absolute percent change in the mean overall fraction of native contacts 〈*Q*_overall_〉 (in this case, computed over all residues in secondary structures within each protein, rather than for individual domains or interfaces) between each protein’s native state simulations and the mean *Q* in the final 10 ns of each misfolded trajectory (Figure 2b). We observe that 76% of misfolded trajectories (1,242 out of 1,631) have ≤20% change in mean *Q* in comparison to the native state, while 58% of trajectories (939 out of 1,631) misfold and have a ≤10% change in *Q*. 9% of trajectories (144 out of 1,631) have a ≤1% change in *Q*. These calculations indicate that a large proportion of trajectories that misfold populate states that are native-like. Therefore, many *E. coli* proteins can populate kinetically trapped near-native conformations that are structurally similar to the native state.

### Misfolded states can persist for days or longer after release from the ribosome

Misfolded conformations that persist for just a few minutes before properly folding are unlikely to have downstream consequences in a cell. To estimate the range of folding times for misfolded conformations we computed the survival probability of the unfolded state, *S*_U_(*t*), for each protein domain and interface and extracted their characteristic folding timescales using a three-state folding model, which reports folding timescales for the fast- and slow-folding phases (see Methods). For a protein to be considered folded all its component domains and interfaces must be folded. Furthermore, since folding pathways that pass through misfolded states take longer to reach the native state, the slow-folding phase reflects the time scale of these pathways. Therefore, for a given protein, we interpret the longest, slow-folding phase time as the time scale of the misfolded state reaching the native state. In total, we are able to reliably determine folding times for 73 out of 122 proteins, with fit equations for other domains having small Pearson *R*^2^ values indicative of low-quality estimates. These extrapolated folding times for the slow phase were then mapped onto experimental times using the acceleration factor associated with the coarse-grained model^28^ (see Methods). The 25^th^, 50^th^, 75^th^, and 95^th^ percentile mean folding times for the slow phase are 1.41 s, 50.9 s, 1.19 x 10^7^ d, and 3.83 x 10^16^ d, respectively, and the full range of times extend from 0.04 s to 1.08 x 10^22^ d (Figure 2c, Table S5). While values at very long times have larger uncertainties, as small differences in the fit parameters will lead to large variation in the extrapolated folding times, these results clearly indicate that many of these misfolded states can persist for many days or longer after synthesis.

### Half of the proteome misfolds and bypasses the chaperone machinery

Misfolded proteins are engaged by various chaperones both co- and post-translationally that help direct their correct folding. So, we next determined how many of the trajectories that exhibit misfolding in our simulations are likely to evade chaperone-dependent quality control mechanisms. As was done for Luciferase, we considered the interactions of each of our 1,631 misfolded trajectories with TF, GroEL/GroES, and DnaK based on the relative difference between the SASA of specific subsets of residues in the misfolded ensemble versus the native state ensemble (see Methods, Eqs. 9–11). We find that 1,053 misfolded trajectories, representing 70 unique proteins, are not likely to interact with TF, as they display 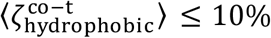 or are too short to engage with it co-translationally (see Methods, Table S6, and Figure S2). A total of 1,411 of misfolded trajectories representing 80 unique proteins are either not known GroEL/GroES^44–46^ clients or have 〈*ζ*_hydrophobic_〉 ≤ 10%, and are therefore not likely to interact excessively with GroEL/GroES (Table S6, Figure 2d). Finally, we find that 1,115 misfolded trajectories representing 74 unique proteins are either not confirmed DnaK clients or have 〈*ζ*_DnaK_〉 ≤ 10% and are therefore unlikely to interact with DnaK excessively (Figure S3). A total of 731 trajectories representing 64 different proteins are misfolded and unlikely to interact with TF, GroEL/GroES, or DnaK (Table S6, Figure S4). These results indicate that 52% of proteins in our representative sample (64 out of 122 unique proteins) exhibit misfolded subpopulations that can bypass chaperones.

### Half of the proteome misfolds and remains soluble

We next assessed how many of the 1,631 trajectories in which the protein misfolds represent conformational states that are likely to remain soluble. For each protein we computed 〈*ζ*_agg_〉, the average relative difference in SASA of residues predicted to be aggregation prone computed over the final 100 ns for each misfolded trajectory, to quantify the difference in aggregation propensity for the misfolded population relative to the native state simulations (see Methods, Eq. 12). Of the 1,631 misfolded trajectories, 814 have 〈*ζ*_agg_〉 ≤ 10%, indicating they are not likely to aggregate in excess of what is observed for the native state (Table S7, Figure S5). We conclude that these trajectories, representing 56% of the proteins in the sample (68 out of 122), are unlikely to aggregate.

### Half of the proteome misfolds and does not exhibit excess degradation

Next, we examined how many misfolded proteins are likely to avoid rapid degradation. We did this by computing 〈*ζ*_hydrophobic_〉, which characterizes the percent difference between the total hydrophobic SASA of misfolded trajectories in comparison to the set of native-state simulations (Eq. 10). The values of 〈*ζ*_hydrophobic_〉 for 896 misfolded trajectories are ≤10%, indicating they are unlikely to be targeted for degradation. These 896 misfolded trajectories predicted to bypass degradation represent 57% (70 out of 122) unique proteins (Table S7, Figure 2d). Thus, a majority of proteins can populate, to varying degrees, misfolded states that are not expected to be degraded at rates much faster than their native fold.

### Half of the proteome misfold into conformations that bypass all aspects of the proteostasis machinery in *E. coli*

Misfolded conformations that do not engage chaperones, do not aggregate, and are not degraded in excess of the native state will remain soluble within the cell for a similar time scale as the native state. We cross-referenced our lists of misfolded trajectories that fall into each of these categories, finding that 8% of all trajectories simulated (517 out of 6,100) misfold into such soluble conformations, and 47% of proteins (57 out of 122) have at least one such trajectory (Table S8). Thus, nearly half of proteins in our sample have subpopulations of misfolded states that will bypass all aspects of protein homeostasis and stay misfolded for biologically long time periods.

### Half of the proteome misfolds and will exhibit altered function

Next, we examined what percentage of the proteome misfolds and is likely to exhibit reduced function. To answer this question, we constructed a database identifying residues that take part in the function of each of the 122 proteins in our data set based on information available in PDB and UniProt database entries. These functional residues were identified based on whether they were in contact with substrates (such as other biomolecules, small-molecule compounds, or ions) in their PDB structures, as well as based on UniProt’s identification of functional residues (see Methods). We then computed the mean relative difference in the structural overlap function of these functional residues in the final 100 ns of misfolded trajectories relative to the native state reference simulations (〈*χ*_func_〉, see Methods). We find that 62% of misfolded trajectories (1,019 out of 1,631) have 〈*χ*_func_,〉 ≥ 10%, indicating that structure at their functional sites are significantly perturbed as well as their function. These trajectories represent misfolded conformations of 69 unique proteins, indicating that 57% of the proteome can populate misfolded conformations likely to exhibit reduced function (Table S8, Figure 2e).

### One-third of proteins exhibit soluble, misfolded, native-like states with reduced functionality

We next determined which of our 122 proteins misfold, evade chaperones, aggregation, degradation, and display reduced function. We find that 31% of proteins (38 out of 122) and 3% of all trajectories (186 out of 6,100) can bypass proteostasis machinery and display decreased function (Table S8, Figure 2f). The extrapolated folding times of these 38 soluble but non-functional proteins range from 2.13 s to 1.07 x 10^22^ days with a median predicted folding time of 1.05 x 10^12^ days, indicating that their function is likely to be perturbed for long timescales.

### Intra-molecular entanglement drives long-lived, soluble misfolded conformations

To determine what characteristics, if any, the misfolded conformations of different proteins have in common, we used the Gauss linking number calculated from linking between a closed loop formed by a native contact between residues *i* and *j* and the pseudo-closed loops formed by the flanking termini, *g*(*i*, *j*)^47^. This quantity provides a useful measure of whether subsections of the protein chain are entangled with each other^48^. Misfolding, or changes in the linkage between the two closed loops, can then be identified by changes in the Gauss linking number of specific native contacts between a reference structure and a target structure (Figure 3a, b).

**Figure 3.**
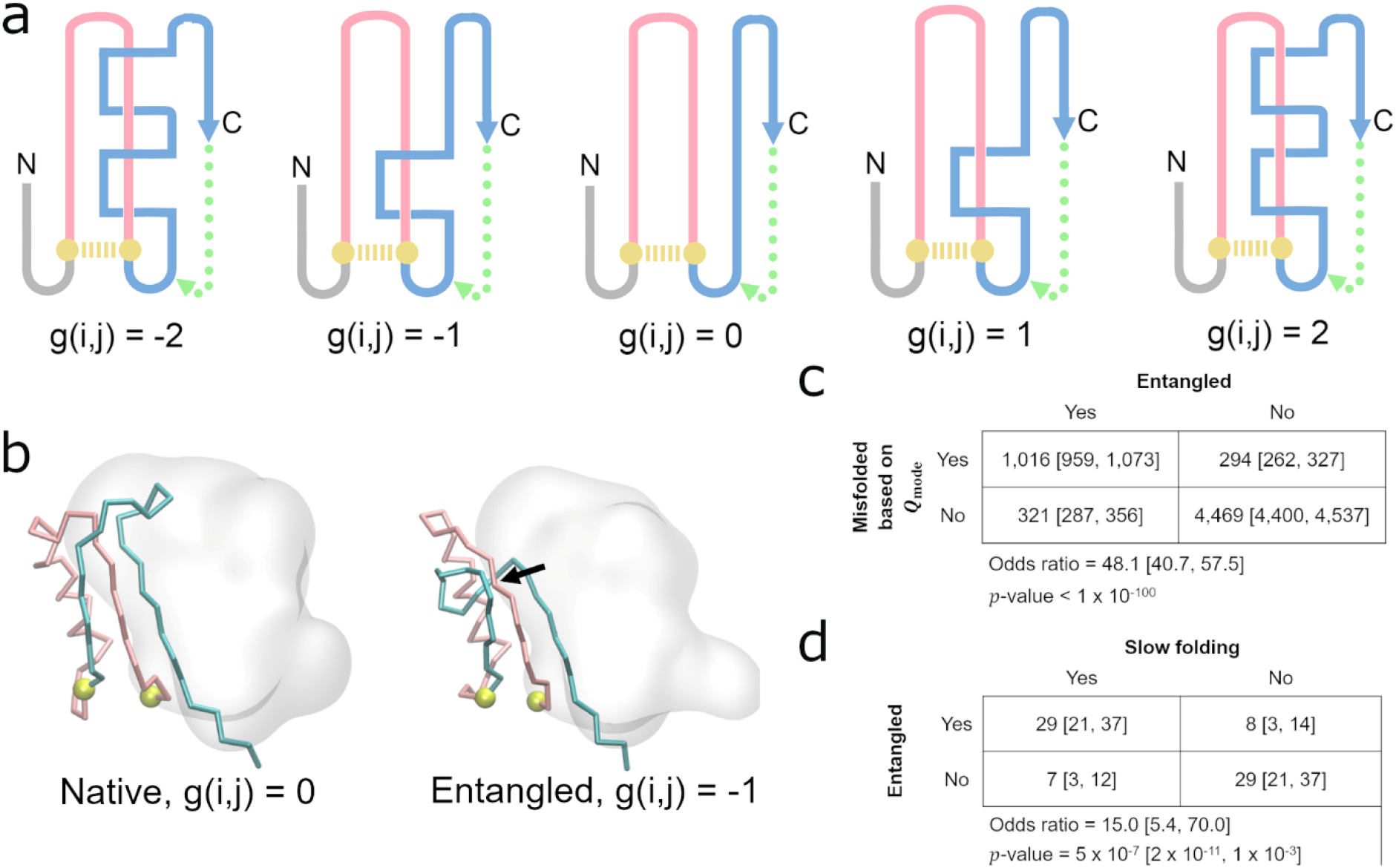
Detecting non-native entanglements in a monomeric protein structure. (a) Schematic of how self-entanglements can be detected by examining the change in the Gauss linking number *g*(*i*, *j*) (Eq. 5) between a closed loop (pink) formed by the backbone segment between residues *i* and *j* that form a native contact (gold dashed line) and another pseudo-closed loop formed by the C-terminal backbone segment (blue) and a pseudo-vector (dashed green line) connecting the C-terminal residue and the start of the C-terminal segment, which begins at residue *j*+1. Threading of the N-terminal segment (composed of residues 1 through *i*-1) is determined in a similar manner. Examples of different Gauss linking numbers and their corresponding structures are shown in this hypothetical illustration. The magnitude of *g*(*i*, *j*) is proportional to the number of threading events of the blue segment through the pink loop, while its sign is a function of the relative positioning of primary structure vectors at crossing points between the pink and blue segments. The structure with *g*(*i*, *j*) =0 exhibits no entanglement. (b) An example of a gain in entanglement of the protein YJGH (PDB: 1PF5), where the C-termini (cyan) threads a loop (pink) formed by the native contact between residues D72 & Y104 (gold). Black arrow indicates the location of the crossing point of the two entangled loops. (c) Contingency table indicating the number of trajectories that are misfolded/folded across our 122 proteins based on *Q*_mode_ analysis and entangled/not entangled. Indicated *p*-values and odds ratios were computed in SciPy using the Fisher Exact Test. (d) Same as (c) except contingency table displays the number of proteins that are entangled/not entangled and predicted to be slow folding/fast folding. For the purposes of this analysis, a protein is considered slow- or fast-folding if its computed folding time is above or below the median folding time from the set of 73 proteins with reliable estimates, respectively. A protein is considered entangled if ≥50% of its misfolded trajectories are entangled. All error bars are 95% confidence intervals computed from bootstrapping 10^6^ times.

To determine whether or not misfolded states tend to be entangled, we generated a 2-by-2 contingency table (Figure 3c) tabulating the co-occurrence of misfolding based on 〈*Q*_mode_〉 and the presence of an entanglement. We find an odds ratio of 48.1 (*p* < 10^−100^, Fisher’s Exact test), indicating that entanglement and misfolding frequently co-occur, with 82% of misfolded states containing an entanglement. Thus, misfolding is predominantly driven by entanglement of segments of the nascent protein with each other.

We hypothesized that due to the large energetic barrier needed to disentangle entanglements, the most long-lived misfolded states in the *E. coli* proteome would tend to be entangled. To test this hypothesis, we generated a second 2-by-2 contingency table and counted how frequently slow- and fast-folding proteins tend to be entangled (Figure 3d). Proteins with an extrapolated folding time for the slow phase greater than the median were considered to be slow folding; and a protein’s misfolded state is considered entangled if ≥50% of its misfolded trajectories display an entanglement. We find an odds ratio of 15.0 (*p* = 5.0 x 10^−7^, Fisher’s Exact test) indicating that the presence of entangled misfolded structures are 15 times more likely to be associated with slow folding. Thus, entanglement is the primary cause of long-lived misfolded states.

Finally, we further hypothesized that, because entangled conformations can represent local minima with only small structural perturbations relative to the native state, the set of 186 trajectories predicted to bypass proteostasis machinery to remain soluble but non-functional for long timescales should be enriched in entangled structures. We find that 94% of these trajectories are entangled (174 out of 186), and that there is a strong association between escaping proteostasis machinery and the presence of an entanglement (odds ratio 59.2, *p* = 3.4 x 10^−102^, Fisher’s Exact test; Figure 4).

**Figure 4.**
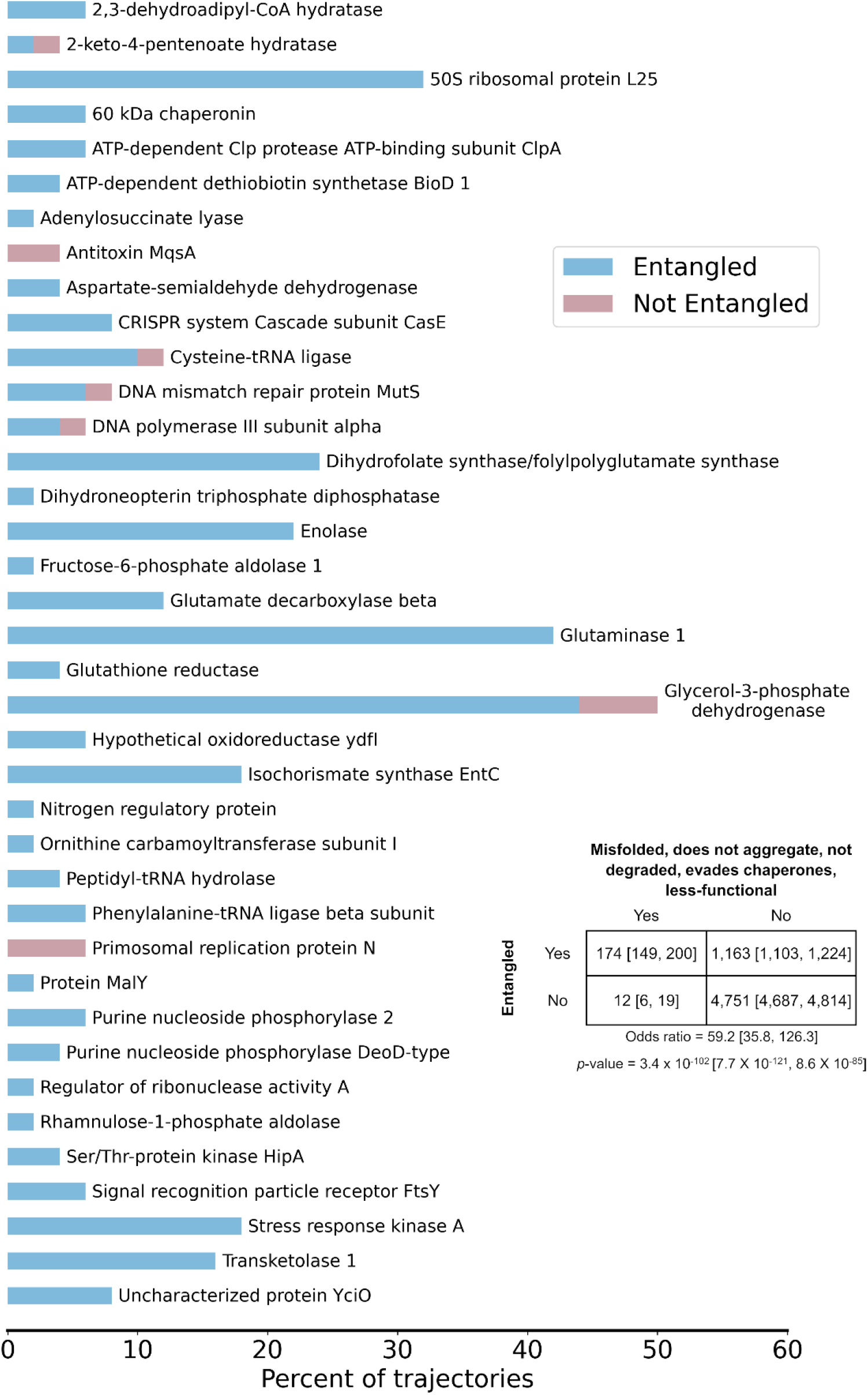
The vast majority of trajectories predicted to bypass cellular quality controls and exhibit reduced function are entangled. The percent of trajectories out of 50 for each of the 38 proteins that bypass quality controls and are predicted to have reduced function that are entangled (blue) or not entangled (red). A total of 174 out of 186 trajectories are entangled. Protein names were taken from UNIPROT; see Table S2 for the structures used and their corresponding gene names. Inset contingency table indicates the number of trajectories that are misfolded and escape proteostasis machinery while remaining non-functional and entangled/not entangled. Indicated p-value and odds ratio were computed in SciPy using the Fisher Exact Test. All error bars are 95% confidence intervals from bootstrapping 10^6^ times.

Taken together, these results demonstrate that the formation of entanglements entangled misfolded states lead to long-lived kinetic traps that can bypass the proteostasis machinery.

### An in-depth case study

To illustrate our key findings, it is useful to consider the structural basis of misfolding for one protein in-depth. We focus on glycerol-3-phosphate dehydrogenase, which has the largest proportion of misfolded trajectories that are predicted to bypass the proteostasis machinery (see Figure 4). It consists of two domains composed of residues 1-387 and 388-501 (Figure 5a). As part of its biological function, glycerol-3-phosophate dehydrogenase uses a flavin adenine dinucleotide cofactor (Figure 5a, dark blue). In our post-translational simulations 74% (=37/50) of trajectories misfold, yet they only exhibit a 4.7% decrease in the fraction of native contacts relative to the native-state simulations (Figures 5b, c). Thus, these misfolded states resemble the native ensemble. This protein also folds extremely slowly, with Domain 1, Domain 2, and the interface between Domains 1 and 2 estimated to require, respectively, on the order of 10^16^, 10^15^, and 10^21^ seconds to fold (Figure 5d). These misfolded states are, however, expected to evade chaperones, aggregation, and degradation to remain soluble based on the similarity of their surface properties to that of the native ensemble (Figure 5e). Twenty-five misfolded trajectories (50%) also exhibit notably reduced structure at functional sites, including around the cofactor, despite being well folded overall (Figure 5f, g, and h). In 92% (=34/37) of these misfolded trajectories a non-native entanglement is present. These results exemplify how entangled misfolded states can perturb portions of a protein critical for function in ways that are structurally subtle compared to gross deformations typically associated with misfolded proteins (Figure 5d and h).

**Figure 5.**
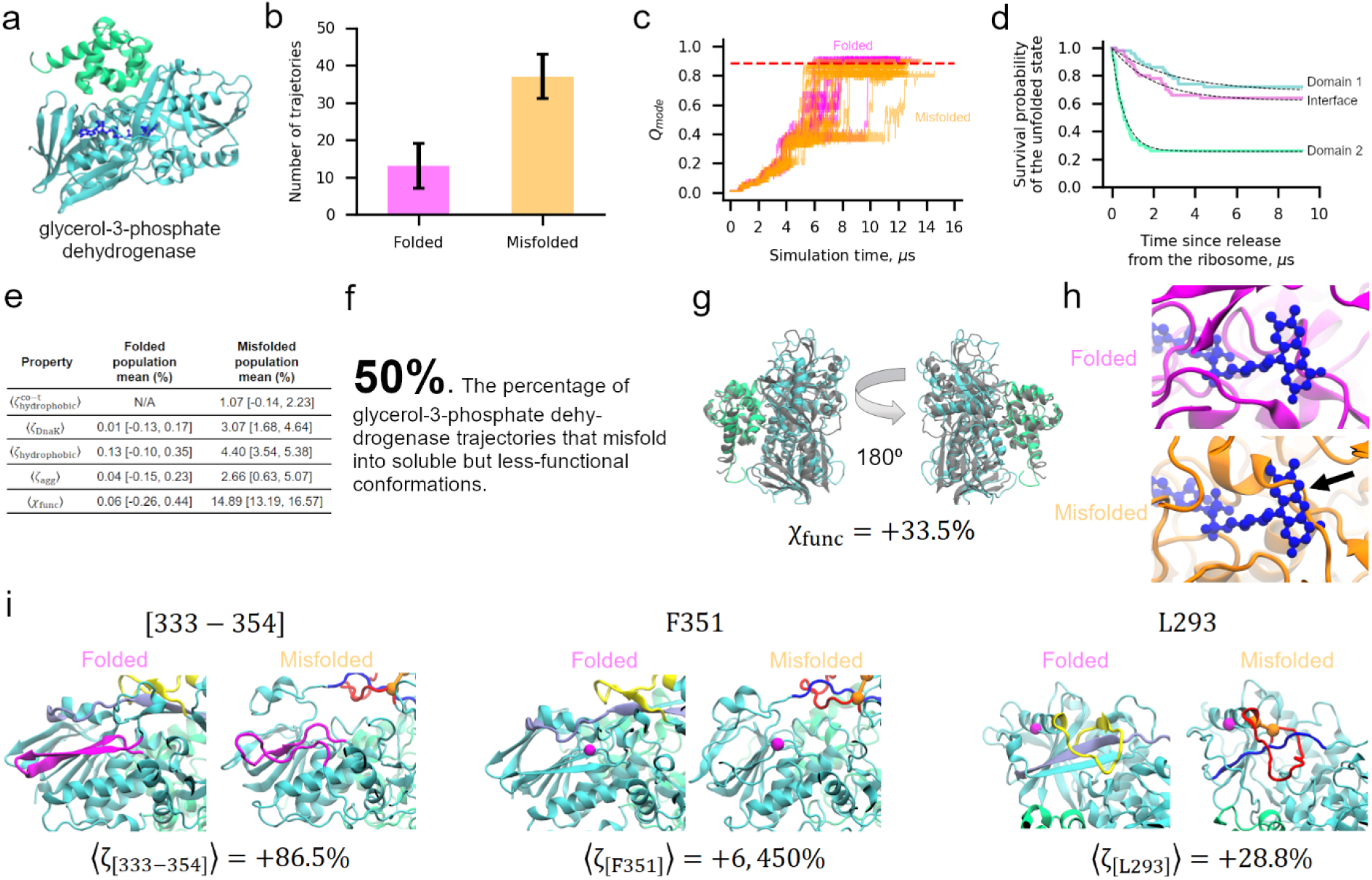
The two-domain glycerol-3-phosphate dehydrogenase protein displays widespread misfolding into soluble but less-functional conformations. (a) Ribbon structure of glycerol-3-phosphate dehydrogenase (PDB ID: 2QCU). Domains 1 and 2 are composed of residues 1-387 and 388-501 and are colored cyan and green, respectively. The FAD cofactor is shown in a dark blue representation. (b) Number of folded and misfolded trajectories for this protein in our simulations. (c) *Q*_mode_ versus time for Domain 1 for the subpopulations of folded (magenta) and misfolded (orange) trajectories. Each line represents one independent trajectory. (d) Survival probability of the unfolded state versus time computed for Domain 1 (cyan), Domain 2 (green), and the Domain 1|2 interface as described in Methods. Dotted black lines are double-exponential fits used to extract rate constants. (e) Summary of key parameters for the folded and misfolded populations. (f) 50% [36%, 64%] of glycerol-3-phosphate dehydrogenase trajectories are predicted to remain soluble but non-functional. (g) Representative misfolded structure colored as in (a) aligned to the native-state reference structure. Despite a high *χ*_func_ value indicative of a less-functional conformation, the protein is largely native. (h) Representative folded and misfolded structures back-mapped to all-atom resolution and then aligned to the native state based on the FAD binding pocket residues. Steric conflict (indicated by black arrow) can be seen between the substrate binding location and the misfolded binding pocket, indicating reduced function of this conformation. (i) Three pairs of structures corresponding to the native state (left) and the first representative structure of metastable state S2 (right) with locations of [333-354], F351, and L293 indicated in magenta. The loop (residues 271-288) and threading (residues 218-237) segments of the entanglement present in S2 are shown in red and blue, respectively. The same regions are colored yellow and light purple in the native state for reference, though no entanglement is present. The CA atoms of residues 271 and 288 that form the contact closing the loop segment are represented by orange spheres. Values of 〈*ζ*_peptide_〉 were calculated with Eq. 16; error bars are available in Table S11. All error bars are 95% confidence intervals computed from bootstrapping 10^6^ times.

### An experimental test for structural changes associated with entanglement

To test these predictions for glycerol-3-phosphate dehydrogenase we carried out protease digestion mass spectrometry (see Methods) in which whole extracts from cells were globally unfolded by incubation in 6 M guanidinium chloride, and refolded by rapid dilution. The structures of the refolding proteins were then interrogated with pulse proteolysis with proteinase K (PK), which specifically cuts at exposed or unstructured sites. The resulting fragments were identified and quantified with mass spectrometry and compared to those from native lysates that were never unfolded. Protease digestion was carried out at 1-min, 5-min, and 120-min timepoints after refolding conditions were established, and glycerol-3-phosphate dehydrogenase’s digestion pattern is observed to change over these time points (see Methods and Supplementary Data File 1). We consider in our analysis only those peptides that show a greater than 3.5-fold difference in abundance in the refolded sample versus native sample 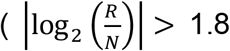, Column W in Supplementary Data File 1 sheet labeled “GlpD”) and whose difference is statistically significant (*p* < 0.01, −log_10_(*p*) > 2, Column Y in same sheet, see Methods^49^). A total of ten unique peptides meet these criteria at one or more experimental time points. At 1 min residue V203 and residues [333-354] are significantly more exposed in the refolded sample than in the native sample. At 5 min L293, F351, Q487, and P387 are more exposed in the refolded than native sample, while V203 and [333-354] are no longer found to be different between refolded and native. After 120 min, seven exposed peptides are found: [333-354], F351, and L293 once again appear more exposed in refolded than native, while Y313, [284-302], D437, and G422 emerge as more exposed in the refolded sample. No one peptide is found to be more exposed in the refolded than native samples at all three time points, though L293, F351, and [333-354] are more exposed at two time points. These experimental data indicate that some glycerol-3-phosphate dehydrogenase molecules that fail to arrive at the native structure rapidly populate misfolded structures.

To test if the entanglements we observe in simulations of glycerol-3-phosphate dehydrogenase can explain these digestion patterns, we structurally clustered the coarse-grain conformations from the final 100 ns of our simulations (based on their *G* and *Q*_overall_) into eight metastable states denoted {S1, S2,…, S8}. We focus on segment [333-354] and residues F351 and L293 because these peptides persist in the experiments, being present at either the 1- or 5-min timepoints *and* the 120-min timepoint. In seven out of eight states an entanglement is present, with states S5, S7, and S8 the most native like (Table S11). The entangled loop or threading segments in these states overlap with one or more peptide fragments in five out of eight states. Structure near cleavage sites is still perturbed even when the entangled region does not overlap with them. For example, the threading of residues 218-237 through the loop formed by residues 271-288 in S2 does not contain residues L293, F351, or segment [333-354]. However, this entanglement increases the solvent accessible surface area of these segments (Figure 5i). This is most clearly seen for F351, which in the native state forms part of a β-sheet buried beneath the threading segment residues (Figure 5i, middle panel, “Folded” structure). When this set of residues becomes entangled by threading through the loop (“Misfolded” structure in Figure 5i), the thread is kinetically trapped in a position that exposes F351 much more than in the native fold. Calculating the solvent accessible surface area change (Eq. 16) of these fragments in each metastable state we find broad agreement with the experimental data (Table S11). Each of the seven entangled metastable states displays increased solvent accessibility at each of the three locations.

## DISCUSSION

Previous work has established that soluble, long-lived, non-functional protein misfolded states can arise from alteration of translation-elongation kinetics. To the best of our knowledge, this study is the first to estimate the extent of this phenomenon across the nascent proteome of an organism and examine the structural and kinetic properties of these kinetically trapped states. We predict that a majority of cytosolic *E. coli* proteins exhibit subpopulations of misfolded, kinetically trapped states, and that many of these misfolded states are similar enough to the native state to evade the proteostasis machinery in *E. coli*. We estimate that one-third of cytosolic *E. coli* proteins have subpopulations that misfold into near-native conformations that have reduced function and bypass the proteostasis network to remain soluble and non-functional for days or longer.

To appreciate these results, it is useful to understand the types of misfolding that can and cannot occur in our simulation model. The coarse-grain forcefield is parameterized for each protein based on its crystal structure, with this native-state conformation encoded as the potential energy minimum in the form of a Gō-based energy function (Eq. 1). This means that the native state is the global free energy minimum at our simulation temperatures; any other state is metastable. Another consequence of this type of model is that misfolding involving non-native tertiary structure formation is not possible. Thus, the misfolded states our model can populate are topologically frustrated states that are kinetic traps^50^. A kinetic trap is a local minimum separated from other conformations in the ensemble by energy barriers much larger than thermal energy, making the attainment of the native state a slow process for some protein subpopulations. In our model, intra-molecular entanglements can occur. These entanglements consist of two parts: a contiguous segment of the protein that forms a ‘closed’ loop, where the loop closure is geometrically defined as a backbone segment that has a native contact between two residues at its ends, and another segment of the protein that threads through this loop (Figure 3).

Proteins can misfold by a variety of mechanisms. For example, Bitran and co-workers^51^ suggest that non-native contacts appear to play an important role in kinetic trapping of some proteins. Misfolding has also been observed via domain swapping^52^, in which highly similar portions of proteins swap with one another. Our results are not mutually exclusive with these other types of misfolding; indeed, one can imagine situations in which domain swapping involves the introduction of an entanglement, or in which non-native contacts form entanglements. Understanding the overlap and interplay of these various types of misfolding in real systems is an interesting open question.

It was previously hypothesized^53^ that like protein topological knots^54^ (which persist when pulling on both termini), this type of entanglement, which we refer to as a non-covalent lasso entanglement^55^, would generate topological frustration and be a kinetic trap. Simulations of proteins with topological knots in their native state^50^ observed that the wrong knot could form and that many of these states were long-lived kinetic traps as they required ‘backtracking’^56^ (*i.e*., unfolding) to fix the knot. While only 3 of the 122 proteins (gene names rlmB, metK, and rsmE) in our study contain topological knots in the native state, the non-covalent lasso entanglement intermediates we observe are non-native pseudoknots^54^ (which unravel when pulling on both termini), and require either reptation of the threaded protein segment out of the closed loop (Figure 3) or local unfolding of the loop surrounding the threaded segment to disentangle. Thus, the results of this study bridge the rich field of polymer topology with biologically important consequences for in vivo protein structure and function.

A potential criticism of this work is that the non-native entanglements we observe may be an artifact of our coarse-grained modeling of proteins. Several lines of evidence indicate this criticism is unfounded. A sufficiently long linear polymer performing a random walk will always sample knotted structures. Thus, it is a fundamental polymer property that knots and entanglements have the potential to form^57^. In a recent study, four entangled structures produced from our coarse-grained model were back-mapped and simulated using classical, all-atom molecular dynamics^32^. The entanglements, and native-like structure of these states persisted for the entire 1 μs simulation time. In another study, one-third of the protein crystal structures in the CATH database were found to contain in the native state the same types of entanglements we observe as intermediates^58^. Taken together, these results indicate that the entanglements we observe are realistic non-native intermediates that have the potential to be populated by many proteins.

Two differences between our simulations and the limited-proteolysis data we compare to lie in the preparation of the proteins and limits of detection. In the experiment proteins are prepared in a chemically denatured state and then allowed to refold, compared to folding concomitant with or after translation. Protein’s that are prone to misfolding during translation are likely to be prone to misfolding during bulk refolding. Thus, while it is not necessary that the same misfolded states be populated under these two different situations, the consistency between the misfolded entangled states of glycerol-3-phosphate dehydrogenase and the persistent and significant protease fragments from the experiment indicates similar misfolded states do occur. Secondly, our computational workflow can detect proteins that misfold as little as 2% of the time (1 misfolded trajectory out of 50); on the other hand, to filter signal from noise, protein regions are only considered more (or less) exposed in the refolded form relative to native if the corresponding PK-fragment is >2-fold (*i.e*., 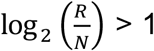 with *p* < 0.01) more (or less) abundant in the proteolysis reaction. Thus, the sensitivity of this approach to misfolded states with low populations is lower, and needs to be considered when comparing to the simulations results.

The majority of proteins simulated in this study misfold in some capacity. And 38 unique proteins, or about one-third of the proteins we simulated, have one or more trajectories that remain soluble and non-functional due to misfolding. Projecting this proportion across the entire set of 2,600 proteins that make up the cytosolic *E. coli* proteome, we estimate that approximately 806 proteins may exhibit misfolding into soluble states. Given that a reduction in the function of a protein has the potential to influence multiple cellular processes, this result suggests that these misfolded states could exert wide-spread influences on cell behavior and phenotype. We also note that it is not only proteins that bypass all aspects of proteostasis and remain non-functional that can negatively impact cells. For example, protein conformations that avoid chaperones and degradation but then go on to aggregate may lead to the accumulation of amyloid fibrils.

Changes to the speed of translation, such as those that may be introduced by synonymous mutations in a protein’s mRNA template, can strongly influence the ability of proteins to fold^7^. The simulation results described here were generated using the wild-type translation rate profile for each protein (see Methods). Experiments have shown that changing translation speed can alter the subpopulation of soluble, less functional states several fold. Thus, the population of proteins that misfold have the potential to be significantly altered in our computer simulations what translation-elongation rates are altered. These complexities make exploration of the influence of translation kinetics on the propensity of the *E. coli* proteome to misfold an interesting direction for future research.

One of the most fundamental timescales of a protein is its half-life, which gives a measure of the lifetime of a typical copy of a protein between its synthesis and degradation. If misfolded states persist on the same timescale as the protein half-life then protein function will be perturbed for most of that protein’s existence. Unfortunately, we are unaware of any proteome-wide studies of protein half-lives in *E. coli*. However, based on studies of protein lifetimes in budding yeast^59^ and human cells^60^, which found median half-lives of 43 min (range 2 min to 81 days) and 36 h (range: 8 h to 153 days), respectively, we estimate that typical half-lives in *E. coli* range from minutes to hours. Many of our extrapolated folding times from our simulations are on the same order of magnitude as these values or greater, indicating that misfolded but states with reduced function can persist for the entire lifetime of a protein. This is consistent with the experimental observation that misfolding can influence folding and function for extended periods^1,2,61^.

If the half-life is a fundamental time scale of a protein, then the cell-division time is a fundamental time scale of a bacterium. In *E. coli*, doubling times during exponential growth phase range from tens of minutes to hours depending on the growth medium^62^. A total of 31 of our 122 proteins have extrapolated folding times for the slow phase longer than 40 min. And, of the 38 proteins that misfold into soluble but less-functional states, 14 have extrapolated folding times longer than 40 min. Since these folding times are on a similar time scale as the doubling time, soluble misfolded conformations will be split between the daughter cells. This suggests that the memory of those events can be encoded in these kinetically trapped states and transferred to the daughter cells. It will be an interesting area of future study to determine whether inheritance of soluble, misfolded proteins with potentially altered function can act as a mechanism for epigenetic inheritance and influence daughter cell behavior.

A key question potentially addressed by our simulations is what allows these misfolded states to remain misfolded in non-functional states for such long timescales? Entanglements appear to allow these misfolded states to persist for long time scales, and their largely native topologies mean they are not excessively acted upon by the proteostasis machinery. In many instances, large-scale unfolding would need to take place in order for the entangled protein to disentangle^50^ to a state from which the native fold is more readily accessible. One interesting avenue for future research is comparison of our results concerning misfolding after ribosomal synthesis with simulations of refolding from denatured chains. Such work would provide a clearer comparison to LiP-MS experiments and enable us to test the hypothesis of whether protein synthesis reduces protein misfolding.

In summary, we have found that the majority of *E. coli* proteins misfold in our simulations, and that some proteins misfold into states that likely bypass cellular proteostasis machinery to remain soluble but with reduced function. We find that these misfolded conformations are able to remain soluble because they are, overall, very similar to the native state, but with certain entanglements that lead to perturbed structure and function. Given that self-entanglement is a fundamental polymer property, the entanglements we have observed represent a universal type of misfolding that has the potential to impact a range of proteins and functions. Specifically, our simulation results suggest the hypothesis that entangled states may be the source of reduced dimerization^2^, enzymatic function^63^, and small-molecular transport^27^ upon changes in translation kinetics induced by synonymous mutations. Future theoretical and experimental efforts should focus on the structural characterization of non-native entangled states and their influence on protein function.

## METHODS

### Selection of proteins and parameterization of their coarse-grain models

A data set of 50 multi- and 72 single-domain proteins was selected at random from a previously developed database of *E. coli* proteins with solved X-ray diffraction or NMR structures^41,64^. This data set contains proteins with realistic distributions of protein size and structural class (see Figure S1 and Table S1, respectively, of Ref. 41). Small sections of missing residues (<10) were rebuilt and minimized in CHARMM, while large missing sections for some multi-domain proteins were rebuilt based on homologous protein structures (see Table S4 of Ref. 41). Each of the rebuilt all-atom models was then converted to a Cα coarse-grain representation. The potential energy forcefield of this coarse-grain model is given by the equation

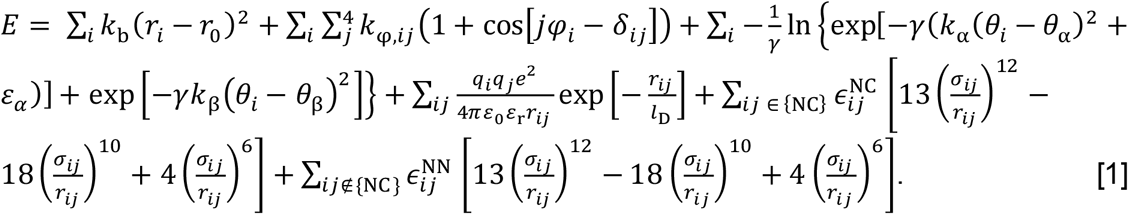

These forcefield terms represent, from left to right, the contributions from Cα – Cα bonds, torsion angles, bond angles, electrostatic interactions, Lennard-Jones-like native interactions, and repulsive non-native interactions to the total potential energy. Full details of the model parameters can be found in Ref. 41. The value of 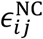, which determines the global energy minimum for a native contact, is calculated as

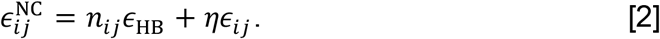

In Eq. 2 *ϵ*_HB_ and *ϵ_ij_* represent energetic contributions from the hydrogen bonds and van der Waals interactions between residues *i* and *j* found within the all-atom structure of the protein, respectively. *n_ij_* is the integer number of hydrogen bonds between residues *i* and *j* and *ϵ*_HB_ = 0.75 kcal/mol. The value of *ϵ_ij_* is initially set based on the Betancourt-Thirumalai potential^65^ and the value of *η* for each individual domain and interface set based on a previously published training set^28^. The values of *η* used for all production simulations are listed in Ref. 41 Tables S2 and S3 alongside all protein names and the chain identifiers used during model building. For simplicity, all proteins are referred to using the PDB ID of the entry from which they were primarily derived. The parameters for the coarse-grain model of firefly luciferase (PDB ID: 4G36) have not previously been reported and are therefore provided in Table S1.

### Simulations of nascent protein synthesis, ejection, and post-translational dynamics

All simulations were performed using CHARMM and the coarse-grain forcefield described in Eq. 1 with an integration timestep of 0.015 ps and a Langevin integrator with friction coefficient of 0.050 ps^−1^ at a temperature of 310 K. The synthesis and ejection of each protein was simulated using a previously published protocol and a coarse-grain cutout of the ribosome exit tunnel and surface (for complete simulation details see Ref. 41). In this model, ribosomal RNA is represented by one interaction site each for each ribose sugar, phosphate group, and pyrimidine base and two interaction sites for each purine bases. Ribosomal proteins are represented at the Cα level. Post-translational dynamics simulations were initiated from the final protein structure obtained after ejection with the ribosome deleted. Fifty statistically independent trajectories were run for each of the 122 proteins in the *E. coli* cytosolic proteome data set and for Luciferase. Post-translational dynamics was run for 30 CPU days for each trajectory. Ten trajectories were also initiated from the native-state coordinates for all proteins and run for 30 CPU days each to provide a realistic reference ensemble for each protein’s folded state.

### Identification of misfolded trajectories

Two order parameters, *Q*_mode_ and *P*(*G_k_* | *PDB*, *traj*), were used to determine whether or a not a given trajectory folds. Detailed definitions of these order parameters are given in the following two Methods sections. A given trajectory is considered to be misfolded if either its *Q*_mode_ or *f_c_*(*G_k_* | *PDB*, *traj*) values (or both, as described below) indicate that the trajectory is significantly different from the native state reference simulations.

### Calculation of *Q*_mode_ and its use as an order parameter for protein folding

The fraction of native contacts, *Q*, was calculated for each domain and interface of all 122 proteins during their synthesis, ejection, and post-translational dynamics. Only contacts between pairs of residues both within secondary structural elements as identified by STRIDE^66^ based on the final rebuilt all-atom structures were considered. To determine when a given domain or interface within a protein folded, the mode of the *Q* values over a 15-ns sliding window (*Q*_mode_) was compared to the representative value of the native state computed as the average *Q*_mode_ over all windows of the ten native-state simulations denoted 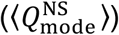. A given trajectory is defined as misfolded if its average *Q*_mode_ over the final 100 ns of the post-translational dynamics portion of the simulation, denoted 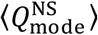, is less than 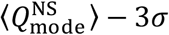, where *σ* is the standard deviation of 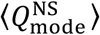.

### Generation of entanglement metric distributions and use as an order parameter for protein folding

To detect non-covalent lasso entanglements we use linking numbers^67^. A link is defined as the entanglement of two closed curves; here, we use (1) the closed curve composed of the backbone trace connecting residues *i* and *j* that form a native contact and (2) the open curves formed by the terminal tails. The native contact between *i* and *j* in (1) is considered to close this loop, even though there is no covalent bond between these two residues. Outside this loop is an N-terminal segment, composed of residues 1 through *i* – 1, and a C-terminal segment, composed of residues *j* + 1 through *N*, whose entanglement through the closed loop we characterize with partial linking numbers denoted *g_N_* and *g_C_*^47^. For a given structure of an *N*-length protein, with a native contact present at residues (*i*, *j*), the coordinates ***R**_l_* and the gradient d***R**_l_* of the point *l* on the curves were calculated as

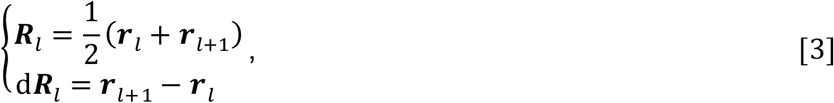

where ***r**_l_* is the coordinates of the Cα atom in residue *l*. The linking numbers *g*_N_(*i*, *j*) and *g_C_*(*i*, *j*) were calculated as

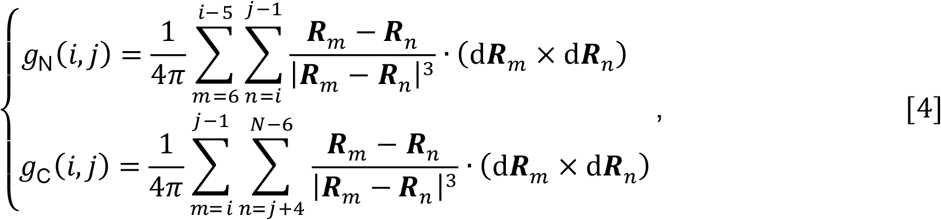

where we excluded the first 5 residues on the N-terminal curve, last 5 residues on the C-terminal curve and 4 residues before and after the native contact for the purpose of eliminating the error introduced by both the high flexibility and contiguity of the termini and trivial entanglements in local structure. The above summations yield two non-integer values, and the total linking number for a native contact (*i*, *j*) was therefore estimated as

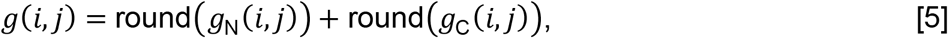

Comparing the absolute value of the total linking number for a native contact (*i*, *j*) to that of a reference state allows us to ascertain a gain or loss of linking between the backbone trace loop and the terminal open curves as well as any switches in chirality. Therefore, there are 6 change in linking cases we should consider (Table S3) when using this approach to quantify entanglement.

To examine the distribution of change in linking (entanglement) detected for a given protein model and statistically independent post-translational trajectory we can generate a discrete probability distribution of the 6 cases in Table S3 as

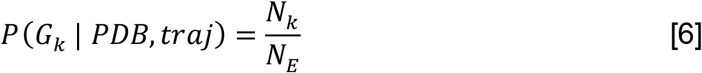

Where *G_k_* for *k* ∈ {0,1,2,3,4,5} is the change in entanglement case of interest from Table S3, *N_E_* is the total number of changes in entanglement instances detected in the trajectory, and *N_k_* is the total number of changes in entanglement in the trajectory of type *k*. As the change in entanglement is held relative to the static crystal structure, it is necessary to correct the post-translational distribution to remove transient changes in entanglement present in the reference state dynamics. This was done by subtraction of the reference distribution from the post-translational distribution considering nontrivial changes in linkage:

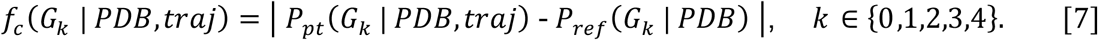

For a given case of entanglement change *k* the magnitude of *f_c_*(*G_k_* | *PDB*, *traj*) increases as the probability of that mode of entanglement change deviates from the reference simulations. We define a given trajectory as misfolded if the average of any of its corrected entanglement values from Eq. 7 over the final 100 ns of the trajectory are ≥0.1.

This trajectory-level analysis is useful for classifying statistically independent sample sets by the level and types of changes in entanglement they exhibit, but a time series metric which conveys the same information was desired to allow for folding time extrapolations. *G* is a time dependent order parameter that reflects the extent of the topological entanglement changes in a given structure compared to the native structure and is calculated as

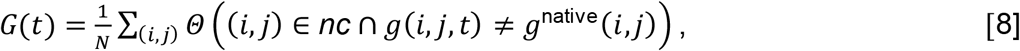

where (*i*, *j*) is one of the native contacts in the native crystal structure; *nc* is the set of native contacts formed in the current structure at time *t*; *g*(*i*, *j*, *t*) and *g*^native^(*i*, *j*) are, respectively, the total entanglement number of the native contact (*i*, *j*) at time *t*, and native structures estimated using Eq. 5; *N* is the total number of native contacts within the native structure and the selection function *Θ* equals 1 when the condition is true and equals 0 when it is false. The larger *G* is, the greater the number of native contact residues that have changed their entanglement status relative to the native state. The utility of entanglement for detecting structural perturbations not apparent by fraction of native contacts or root mean square deviation is visually described in Figure S6.

### Calculation and extrapolation of folding times

Folding times were determined for each domain and interface from their post-translational *Q*_mode_ and *G* time series. The folding time for a domain or interface is taken as the first *t* at which *Q*_mode_ is greater than or equal to 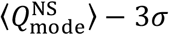 and with *G* ≤ *G_xs_*. The survival probability of the unfolded state was then computed based on these folding times and fit to the double exponential function *S_U_*(*t*) = *f*_1_exp(*k*_1_ * *t*) + *f*_2_exp (*k*_2_ * *t*) with *f*_1_ + *f*_2_ ≡ 1. This double-exponential fit equation represents a kinetic scheme in which the unfolded and misfolded states each proceed irreversibly to the folded state by parallel folding pathways and there is no inter-transitions between unfolded and misfolded states. Folding times for each kinetic phase were computed as 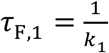 and 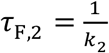, with the overall folding time of the domain or interface taken as the longer of the two folding times. The folding time reported for each protein is the longest folding time from any of its constituent domains or interfaces. The assumption of double-exponential folding kinetics is not a good assumption for all 122 proteins simulated. We therefore only consider domains and interfaces whose *S_U_*(*t*) time series are fit with a Pearson *R*^2^ > 0.90. We are also unable to compute folding times for the nine proteins for which no trajectories folded. In total, we found reliable folding times for 73 of our 122 proteins. Simulated folding times were extrapolated to experimental timescales using the equation *τ*_exp_ = *τ*_sim_ * *α*, where *τ*_sim_ is a given protein’s simulated folding time and α = 3,967,486 is the mean acceleration of folding in our coarse-grain simulations relative to real timescales^28^.

### Identifying misfolded proteins unlikely to interact with trigger factor

To determine whether or not a given protein in our data set is likely to interact with trigger factor (TF) during synthesis we computed the relative difference in the hydrophobic SASA between misfolded trajectories and folded trajectories on the ribosome using the equation

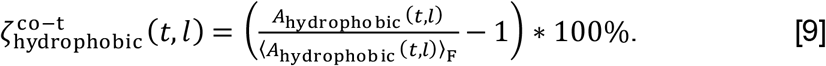

In Eq. 9, *A*_hydrophobic_(*t*, *l*) is the total hydrophobic SASA of residues in the nascent protein at time *t* and nascent chain length *l* exposed outside of the ribosome exit tunnel (defined as having an x-coordinate ≥100 Å in the internal CHARMM coordinate system; see Ref. 41 for details). The term 〈*A*_hydrophobic_(*t*, *l*)〉_F_ is the mean total hydrophobic SASA of residues at time *t* and length *l* outside of the exit tunnel calculated over all frames of synthesis trajectories identified to be folded by *Q*_mode_ and *G* analysis. Note well, the reference states for Equations 10–12 are the native-state reference simulations initiated in bulk solution, but due to the co-translational nature of TF interactions we use the folded subpopulation of synthesis trajectories as the reference state in Eq. 9. As TF is thought to only interact with nascent proteins of 100 residues or longer^68^, we compute Eq. 9 for *l* = {100, 101,…,*N*} for each protein and trajectory, where *N* is the total number of residues in the full-length protein. Proteins shorter than 100 amino acids are considered not to interact with TF. To quantify the overall propensity of a misfolded trajectory of a given protein to interact with TF we compute 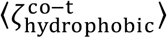 as the mean of 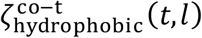 for all *t* and allowed values of *l*. Based on examination of 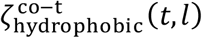 time series, we determined that a value of 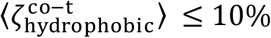 corresponds to proteins that are unlikely to engage TF significantly more than folded conformations on the ribosome. For the purposes of this calculation and all others in this work that consider sets of hydrophobic residues, coarse-grain interactions sites representing {Ile, Val, Leu, Phe, Cys, Met, Ala, Gly, Trp} are considered to be hydrophobic.

### Identifying misfolded proteins unlikely to interact with GroEL/GroES

Our data set of 122 *E. coli* proteins was first cross-referenced with the list of 276 confirmed GroEL/GroES substrates^44–46^. The 103 proteins that do not appear in this list of confirmed clients are considered to not be GroEL/GroES client proteins. GroEL/GroES is thought to identify and bind regions of exposed hydrophobic surface area on nascent proteins. To determine whether the misfolded conformations of proteins that can interact with GroEL/GroES are likely to do so, we compared the total hydrophobic SASA of misfolded conformations with the native-state ensemble using the equation

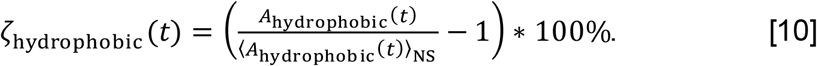

This equation provides the quantity *ζ*_hydrophobic_(*t*), which measures the relative difference between the SASA of hydrophobic residues within a misfolded conformation at time *t* (*A*_hydrophobic_(*t*)) in comparison to to the mean hydrophobic SASA calculated over all frames of the native-state reference simulations (〈*A*_hydrophobic_(*t*)〉_NS_). As GroEL/GroES interacts with proteins post-translationally, Eq. 10 was applied to the post-translational simulation data for all misfolded conformations of a given protein and the average over the final 100 ns computed for each trajectory (〈*ζ*_hydrophobic_〉). The full set of proteins with misfolded conformations that we predict will not interact with GroEL/GroES is taken as the union of the sets of all trajectories for proteins not in the list of experimentally confirmed clients and the list of trajectories with 〈*ζ*_hydrophobic_〉 ≤ 10%.

### Identifying misfolded proteins unlikely to interact with DnaK

We first cross-referenced our list of 122 simulated proteins with a list of DnaK client proteins. The 34 proteins that do not appear on this list are considered to not interact with DnaK. To determine whether misfolded conformations of proteins that can interact with DnaK are likely to do so, we predicted DnaK binding sites using the Limbo webserver^35^ and then compared the total SASA of residues in predicted binding sites between misfolded conformations and the native state using the equation

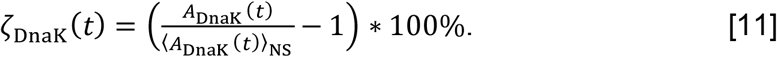

In this equation *A*_DnaK_(*t*) is the total SASA of residues in DnaK binding sites as a function of time *t* and 〈*A*_DnaK_(*t*)〉_NS_ is the mean total SASA of residues in DnaK binding sites averaged over all frames of the native-state reference simulations. Equation 11 was applied to the post-translational simulation data of all misfolded conformations for each misfolded trajectory and the average over the final 100 ns of each trajectory computed (〈*ζ*_DnaK_〉). The full set of trajectories with misfolded conformations that we predict will not interact with DnaK is taken as the union of the sets of trajectories for proteins not in the list of experimentally confirmed clients and the list of trajectories with 〈*ζ*_DnaK_〉 ≤ 10%. We note that three proteins, PDB IDs 2JRX, 2V81, and 2KFW, are predicted by Limbo to have no DnaK binding sites. 2JRX and 2KFW appear on the list of confirmed DnaK clients, and are therefore considered to be DnaK binders. 2V81, however, is not a confirmed DnaK client and is thus counted as not likely to interact with DnaK.

### Identifying misfolded proteins unlikely to aggregate

We used the AMYLPRED2 webserver^37^ to predict the sets of aggregation-prone residues within the primary sequences of our 122 proteins. Whether or not a given trajectory for a protein is likely to aggregate was determined by comparing the SASA of aggregation-prone regions within the misfolded trajectory to the mean SASA of aggregation-protein regions in the native-state reference ensemble using the equation

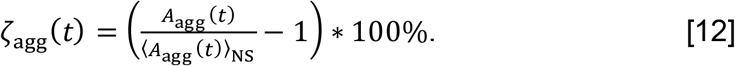

In this equation, *A*_agg_(*t*) is the total SASA of residues in aggregation-prone regions as a function of time, *t*, and 〈*A*_agg_(*t*)〉_NS_ is the mean total SASA of residues in aggregation-prone regions averaged over all frames of the native-state reference simulations. Equation 12 was applied to the post-translational simulation data for all misfolded trajectories of a given protein and the average over the final 100 ns for each trajectory (〈*ζ*_agg_〉). Trajectories with 〈*ζ*_agg_〉 ≤ 10% are considered to be unlikely to aggregate. Note that our methods do not account for the presence of forms of aggregates other than amyloid, as our calculation is based on the AMYLPRED2 algorithm.

### Identifying misfolded proteins unlikely to be degraded

Whether or not a protein’s misfolded conformations are likely to be targeted for degradation was determined on the basis of *ζ*_hydrophobic_ (Eq. 10). Proteins with 〈*ζ*_hydrophobic_〉 ≤ 10% are considered to be unlikely to be degraded.

### Creating a database of functional residues for *E. coli* proteins

Information from UniProt and RCSB was unified and parsed to create a database of residues implicated in function for each of our 122 proteins. Residues involved in interactions with small molecules were identified as those residues with heavy atoms within 4.5 Å of any heteroatom (identified by the HETATM keyword in PDB records) other than water and non-native amino acids such as selenidomethionine (*i.e*., MSE residues). Many proteins must form multimeric complexes in order to exercise their function. To consider these interactions, we also identified residues with heavy atoms within 4.5 Å of heavy atoms in a different chain ID within the same PDB structure.

Our 122 coarse-grain models were built from single PDB structures or, in the case of some multi-domain protein models, the merging of multiple structures. These structures used for model building often lack ligands or protein binding partners that may be required for function due to differences in crystallographic conditions and/or the intention of the original crystallographers. To provide a broader view of functional residues we therefore also considered all PDB structures identified by UniProt to represent the same gene product. Functional residues were identified in these alternative structures as described above for the initial structures. PDB entries representing the same protein often have different residue numbering schemes and a small number of mutations. We therefore used amino acid alignments in BLAST to determine the mapping from alternative numbering schemes to the numbering scheme within the structure used for model building. Only those domains with at least 97% sequence identity were considered for this analysis to allow for small mutational changes while excluding significantly different proteins. A summary of each of the terms in the database and their meanings is provided in Table S10.

### Determining which misfolded conformations are likely non-functional

The relative difference in function between misfolded trajectories and native-state reference trajectories was determined by calculating the relative difference of the structural overlap of residues identified to be involved in protein function using the equation

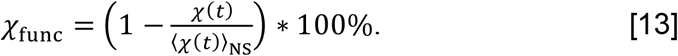

In this equation, *χ*(*t*) is the structural overlap between residues implicated in function at time *t* in a misfolded trajectory with the native-state reference structure and 〈*χ*(*t*)〉_NS_ is the mean of this same value computed over all simulation frames of the native-state reference simulations. The value of *χ*(*t*) is calculated as

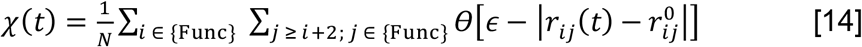

and gives the fraction of pairwise distances that are at native-like values at time *t*. The set {Func} contains all residues implicated in protein function. The indices *i* and *j* correspond to residues in {Func} for the protein being analyzed. *N* is the total number of pairwise contacts between residues *i* and *j* both in {Func} that also satisfy the condition *j* ≥ *i* + 2. The parameters *r_ij_*(*t*) and 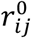 are the distances between residues *i* and *j* at time *t* and between *i* and *j* in the native state reference structure, respectively. *θ*(*x*) is the step function given by

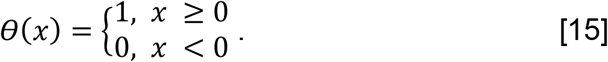

The value of *ϵ* is taken as 0.2 · r_c_α__ where r_c_α__ = 3.81 Å is the virtual bond length between coarse-grain interaction sites in the coarse-grain simulation model. A particular pair of residues *i* and *j* contribute 1 to *χ*(*t*) if 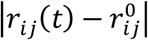 is less than *ϵ*, such that 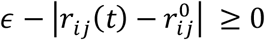, and in all other situations contribute 0 to *χ*(*t*). Similar forms of this equation have been used previously to observe structural transitions in simulations of proteins^69^. The value of *χ*_func_ was averaged over the final 100 ns for given protein trajectory to give 〈*χ*_func_〉, the mean relative difference in structure of functional residues for the misfolded conformation in comparison to the native state. Trajectories with 〈*χ*_func_〉 ≥ 10% are considered to be less functional than the native state.

### Selection of 10% threshold for classifying misfolded trajectories as native-like

We selected a threshold of 10% for *ζ*_hydrophobic_(*t*), *ζ*_DnaK_(*t*), *ζ*_agg_(*t*), and *χ*_func_ (see Eqs. 10, 11, 12, and 13) based on an analysis of how frequently our native state reference trajectories for each protein explore conformations with 10% or greater difference from their native state mean (Table S9). Virtually all proteins explore conformations with ≥10% for each metric in their native state, indicating that 10% is a parsimonious threshold for determining if misfolded conformations are native like. In the case of 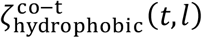, similar calculations as those performed for the other metrics would require running prohibitively expensive arrested-ribosome-nascent chain complex simulations for each of our 122 proteins. We therefore use a threshold of 10% for 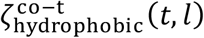 for the sake of consistency with our other thresholds.

### Protease mass spectrometry experiments

See the Supplementary Methods for a full description. Briefly, *E. coli* K12 cells (NEB) were grown in 2 sets of 3 × 50 mL (biological triplicates) MOPS EZ rich media from saturated overnight cultures with a starting OD_600_ of 0.05. As described in Ref. 71, one set was supplemented with 0.5 mM [^13^C_6_]L-Arginine and 0.4 mM [^13^C_6_]L-Lysine and the other with 0.5 mM L-Arginine and 0.4 mM L-Lysine. Cells were cultured at 37°C with agitation (220 rpm) to a final OD_600_ of 0.8. Each heavy/light pair was pooled together; cells were collected by centrifugation at 4000 *g* for 15 mins at 4°C, supernatants were removed, and cell pellets were stored at −20°C until further use.

Frozen cell pellets were resuspended in a lysis buffer consisting of 900 μL of Tris pH 8.2 (20 mM Tris pH 8.2, 100 mM NaCl, 2 mM MgCl2 and supplemented with DNase I to a final concentration (f.c.) of 0.1 mg mL^−1^). Resuspended cells were cryogenically pulverized with a freezer mill (SPEX Sample Prep). Lysates were then clarified at 16000 *g* for 15 min at 4 °C to remove insoluble cell debris. To deplete ribosome particles, clarified lysates were ultracentrifuged at 33,300 rpm at 4 °C for 90 min using a SW55 Ti rotor. Protein concentrations of clarified lysates were determined using the bicinchoninic acid assay (Rapid Gold BCA Assay, Pierce) and diluted to 3.3 mg mL^−1^ using lysis buffer.

To prepare native samples, 3.5 μL of normalized lysates were diluted with 96.5 μL of Tris native dilution buffer (20 mM Tris pH 8.2, 100 mM NaCl, 10.288 mM MgCl_2_, 10.36 mM KCl, 2.07 mM ATP, 1.04 mM DTT, 62 mM GdmCl) to a final protein concentration of 0.115 mg mL^−1^. Native samples were then equilibrated by incubating for 90 min at room temperature. To prepare unfolded samples, 600 μL of normalized lysates, 100 mg of solid GdmCl, and 2.4 μL of a freshly prepared 700 mM DTT stock solution were combined, and solvent was removed using a vacufuge plus to a final volume of 170 μL. Unfolded lysates were incubated overnight at room temperature. To refold, 99 μL of refolding dilution buffer (19.5 mM Tris pH 8.2, 97.5 mM NaCl, 10.03 mM MgCl2, 10.1 mM KCl, 2.02 mM ATP and .909 mM DTT) were rapidly added to 1 μL of unfolded extract. Refolded samples were then incubated at room temperature for 1 min, 5 min or 2 h.

100 μL of the native or refolded lysates was added to Proteinase K (enzyme:substrate ratio of 1:100 w/w ratio^70^), incubated for 1 min at room temperature, and quenched by boiling in a mineral oil bath at 110°C for 5. Boiled samples were transferred to tubes containing 76 mg urea. To prepare samples for mass spectrometry, dithiothreitol was added to a final concentration of 10 mM and samples were incubated at 37°C for 30 minutes. Iodoacetamide was added to a final concentration of 40 mM and samples were incubated at room temperature in the dark for 45 minutes. LysC was added to a 1:100 enzyme:substrate (w/w) ratio and samples were incubated at 37°C for 2 h, urea was diluted to 2 M using 100 mM ammonium bicarbonate pH 8, then trypsin was added to a 1:50 enzyme:substrate (w/w) ratio and incubated overnight at 25°C.

Peptides were acidified, desalted with Sep-Pak C18 1 cc Vac Cartridges, dried down, and resuspend in 0.1% formic acid, as previously described^71^. LC-MS/MS acquisition was conducted on a Thermo Ultimate3000 UHPLC system with an Acclaim Pepmap RSLC C18 column (75 μm × 25 cm, 2 μm, 100 Å) in line with a Thermo Q-Exctive HF-X Orbitrap, identically as previously described^71^.

Proteome Discoverer (PD) Software Suite (v2.4, Thermo Fisher) and the Minora Algorithm were used to analyze mass spectra and perform Label Free Quantification (LFQ) of detected peptides. Default settings for all analysis nodes were used except where specified. The data were searched against Escherichia coli (UP000000625, Uniprot) reference proteome database. For peptide identification, the PD MSFragger node was used, using a semi-tryptic search allowing up to 2 missed cleavages^72^. A precursor mass tolerance of 10 ppm was used for the MS1 level, and a fragment ion tolerance was set to 0.02 Da at the MS2 level. Additionally, a maximum charge state for theoretical fragments was set at 2. Oxidation of methionine and acetylation of the N-terminus were allowed as dynamic modifications while carbamidomethylation on cysteines was set as a static modification. Heavy isotope labeling (^13^C_6_) of Arginine and Lysine were allowed as dynamic modifications. The Philosopher PD node was used for FDR validation. Raw normalized extracted ion intensity data for the identified peptides were exported from the .pdResult file using a three-level hierarchy (protein > peptide group > consensus feature). These data were further processed utilizing custom Python analyzer scripts (available on GitHub, and described in depth previously^49,71^).

### Clustering long-lived misfolded states of glycerol-3-phosphate dehydrogenase

The structural distribution from the last 100 ns of the post-translational simulations of glycerol-3-phosphate dehydrogenase was assessed as the pseudo free energy −*ln*(*P*), where *P* is the probability density, along the order parameters *G* and *Q*_overall_. To further analyze the post-translational structures, 400 clusters (micro-states) were grouped from the last 100 ns trajectories using the k-means algorithm^73,74^. A Markov state model (MSM) was built and the clusters were coarse-grained into a small number of metastable states using the PCCA+ algorithm^75^. The number of metastable states was chosen based on the existence of a gap in the eigenvalue spectrum of the transition probability matrix^76^. Five representative structures of each metastable state were randomly sampled from all microstates according to the probability distribution of the microstates within the given metastable state. All the clustering and MSM building were performed by using PyEmma package^77^.

### Determining which coarse-grain structures have increased exposure of peptides

The relative change in solvent accessible surface area of experimentally identified peptides for glycerol-3-phosphate dehydrogenase was calculated as

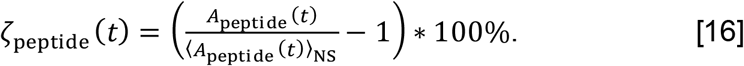

In this equation, *A*_peptide_(*t*) is the total SASA of the residues within the peptide under consideration at time *t* and 〈*A*_peptide_(*t*)〉_NS_ is the mean value of *A*_peptide_(*t*) computed over all frames of the native-state reference simulations. Eq. 16 was applied to all frames in the final 100 ns of each glycerol-3-phosphate dehydrogenase trajectory and then averaged separately for each of its eight misfolded metastable states. Values of 〈*χ*_[333–354]_〉, 〈*ζ*_[F351]_〉, 〈*ζ*_[L293]_〉 are summarized in Table S11 for metastable states {S1, S2,…, S8}.

## DATA AVAILABILITY

Raw data for Figures 1, 2, 3, 4, 5, S1, S2, and S3 are available in the included software directory. We cannot feasibly provide all ~30 TB of trajectory data, but we do provide sample trajectory files and use them to demonstrate our analysis methods. Supplementary Data File 1 contains the annotated experimental data for glycerol-3-phosphate dehydrogenase. Supplementary Data Files 2, 3, and 4 contain all peptide-level data from the 1-, 5-, and 120-min time points, respectively. The mass spectrometry proteomics data have been deposited to the ProteomeXchange Consortium via the PRIDE partner repository with the dataset identifier PXD031425.

## CODE AVAILABILITY

CHARMM input files, Python analysis code, sample commands, and example output are available in the included software directory. A CHARMM license is required to run the molecular dynamics simulation programs. Additional software is available at: https://github.com/obrien-lab/ and https://github.com/FriedLabJHU/Refoldibility-Tools/.

## ACKNOWLEDGMENTS

SDF acknowledges support from the NIH Director’s New Innovator Award (DP2GM140926) and from the National Science Foundation Division of Molecular and Cellular Biology (MCB-2045844). EPO acknowledges funding from the National Institutes of Health (R35-GM124818) and the National Science Foundation (MCB-1553291).

## Supplementary Information

### Supplementary Methods

#### Preparation of K12 Cell Pellets

*E. coli* K12 cells (NEB) were grown in 2 sets of 3 × 50 mL (biological triplicates) of in-house prepared MOPS EZ rich media(-Arginine/-Lysine) from saturated overnight cultures with a starting OD_600_ of 0.05. Similar to reported elsewhere^71^, one set was supplemented with 0.5 mM [^13^C_6_]L-Arginine and 0.4 mM [^13^C_6_]L-Lysine and the other with 0.5 mM L-Arginine and 0.4 mM L-Lysine. Cells were cultured at 37°C with agitation (220 rpm) to a final OD_600_ of 0.8. Each heavy/light pair was pooled together and then transferred to 2 × 50 mL falcon tubes and collected by centrifugation at 4000 *g* for 15 mins at 4°C. The supernatants were removed, and cell pellets were stored at −20°C until further use.

Frozen cell pellets were resuspended in a lysis buffer consisting of 900 μL of Tris pH 8.2 (20 mM Tris pH 8.2, 100 mM NaCl, 2 mM MgCl2 and supplemented with DNase I to a final concentration (f.c.) of 0.1 mg mL^−1^). Resuspended cells were flash frozen by slow drip over liquid nitrogen and cryogenically pulverized with a freezer mill (SPEX Sample Prep) over 8 cycles consisting of 1 min of grinding (9 Hz), and 1 min of cooling. Pulverized lysates were transferred to 50 mL centrifuge tubes and thawed at room temperature for 20 min. Lysates were then transferred to fresh 1.5 mL microfuge tubes and clarified at 16000 *g* for 15 min at 4 °C to remove insoluble cell debris. To deplete ribosome particles, clarified lysates were transferred to 3 mL *konical* tubes and ultracentrifuged at 33,300 rpm at 4 °C for 90 min without sucrose cushions using a SW55 Ti rotor. Protein concentrations of clarified lysates were determined using the bicinchoninic acid assay (Rapid Gold BCA Assay, Pierce) in a microtiter format with a plate reader (Molecular Devices iD3) using BSA as a calibration standard. Protein concentrations were diluted to a standard concentration of 3.3 mg mL^−1^ using Tris lysis buffer. This generates the normalized lysates for all downstream workflows.

#### Preparation of Native and Refolded Lysates for Limited Proteolysis Mass Spectrometry

To prepare half-isotopically-labeled native samples, 3.5 μL of normalized lysates derived from pellets in which half of the cells were grown with [^13^C_6_]L-Arginine and [^13^C_6_]L-Lysine during cell culture and half of the cells were grown with natural abundance L-Arginine and L-Lysine during cell culture, were diluted with 96.5 μL of Tris native dilution buffer (20 mM Tris pH 8.2, 100 mM NaCl, 10.288 mM MgCl2, 10.36 mM KCl, 2.07 mM ATP, 1.04 mM DTT, 62 mM GdmCl) to a final protein concentration of 0.115 mg mL^−1^. Following dilution, the final concentrations are 20 mM Tris pH 8.2, 100 mM NaCl, 10 mM MgCl2, 10 mM KCl, 2mM ATP, 1 mM DTT and 60 mM GdmCl. Native samples were then equilibrated by incubating for 90 min at room temperature prior to limited proteolysis.

The refolding samples were prepared as described previously^49^. Briefly: 600 μL of normalized lysates, 100 mg of solid GdmCl, and 2.4 μL of a freshly prepared 700 mM DTT stock solution were added to a fresh 1.5 mL microfuge tube, and solvent was removed using a vacufuge plus to a final volume of 170 μL, such that the final concentrations of all components were 11.6 mg mL^−1^ protein, 6 M GdmCl, 70 mM Tris pH 8.2, 350 mM NaCl, 7 mM MgCl2, and 10 mM DTT. Unfolded lysates were incubated overnight at room temperature to complete unfolding prior to refolding.

To prepare refolding samples, 99 μL of refolding dilution buffer (19.5 mM Tris pH 8.2, 97.5 mM NaCl, 10.03 mM MgCl2, 10.1 mM KCl, 2.02 mM ATP and .909 mM DTT) were added to a fresh 1.5 mL microfuge tube. 1 μL of unfolded extract was then added to the tube containing the refolding dilution buffer and quickly mixed by rapid vortexing, diluting the sample by 100x, followed by flash centrifugation to collect liquids to the bottom of the tube. The final concentrations were 20 mM Tris pH 8.2, 100 mM NaCl, 10 mM MgCl2, 10 mM KCl, 2mM ATP, 1 mM DTT and 60 mM GdmCl. Refolded samples were then incubated at room temperature for 1min, 5 min or 2 h to allow for proteins to refold prior to limited proteolysis.

To perform limited proteolysis, 2 μL of a PK stock (prepared as a 0.067 mg mL^−1^ PK in a 1:1 mixture of Tris lysis buffer and 20% glycerol, stored at −20°C and thawed at most only once) were added to a fresh 1.5 mL microfuge tube. After refolded proteins were allowed to refold for the specified amount of time (1 min, 5 min, or 2 h), or native proteins were allowed their 90 min equilibration, 100 μL of the native/refolded lysates were added to the PK-containing microfuge tube and quickly mixed by rapid vortexing (enzyme:substrate ratio is a 1:100 w/w ratio^70^), followed by flash centrifugation to collect liquids to the bottom of the tube. Samples were incubated for exactly 1 min at room temperature before transferring them to a mineral oil bath preequilibrated at 110°C for 5 min to quench PK activity. Boiled samples were then flash centrifuged (to collect condensation on the sides of the tube), and transferred to fresh 1.5 mL microfuge tube containing 76 mg urea such that the final urea concentration was 8 M and the final volume was 158 μL. They are then vortexed to dissolve the urea to unfold all proteins and quench any further enzyme activity indefinitely, and flash centrifuged to collect liquids to the bottom of the tubes.

All protein samples were prepared for mass spectrometry as follows: 2.25 μL of a freshly prepared 700 mM stock of DTT were added to each sample-containing microfuge tube to a final concentration of 10 mM. Samples were incubated at 37°C for 30 minutes at 700 rpm on a thermomixer to reduce cysteine residues. 9 μL of a freshly prepared 700 mM stock of iodoacetamide (IAA) were then added to a final concentration of 40 mM, and samples were incubated at room temperature in the dark for 45 minutes to alkylate reduced cysteine residues. 1 μL of a 0.1 μg μL^−1^ stock of LysC (NEB) was added to the samples (to a final enzyme:substrate ratio of 1:100 w/w) and incubated for 2 h at 37°C at 700 rpm. 471 μL of 100 mM ammonium bicarbonate (pH 8) were added to the samples to dilute the urea to a final concentration of 2 M. 2 μL of a 0.1 μg μL^−1^ stock of Trypsin (NEB) were added to the samples (to a final enzyme:substrate ratio of 1:50 w/w) and incubated overnight (15-16 h) at 25°C at 700 rpm (not 37°C, so as to minimize decomposition of urea and carbamylation of lysines).

#### Desalting of Mass Spectrometry Samples

Peptides were desalted with Sep-Pak C18 1 cc Vac Cartridges (Waters) over a vacuum manifold. Tryptic digests were first acidified by addition of 16.6 μL trifluoroacetic acid (TFA, Acros) to a final concentration of 1% (vol/vol). Cartridges were first conditioned (1 mL 80% ACN, 0.5% TFA) and equilibrated (4 x 1 mL 0.5% TFA) before loading the sample slowly under a diminished vacuum (ca. 1 mL/min). The columns were then washed (4 x 1 mL 0.5% TFA), and peptides were eluted by addition of 1 mL elution buffer (80% ACN, 0.5% TFA). During elution, vacuum cartridges were suspended above 15 mL conical tubes, placed in a swing-bucket rotor (Eppendorf 5910R), and spun for 3 min at 350 g. Eluted peptides were transferred from Falcon tubes back into microfuge tubes and dried using a vacuum centrifuge (Eppendorf Vacufuge). Dried peptides were stored at −80°C until analysis. For analysis, samples were vigorously resuspended in 0.1% FA in Optima water (ThermoFisher) to a final concentration of 0.5 mg mL^−1^.

#### LC-MS/MS Acquisition

Chromatographic separation of digests were carried out on a Thermo UltiMate3000 UHPLC system with an Acclaim Pepmap RSLC, C18, 75 μm × 25 cm, 2 μm, 100 Å column. Approximately, 1 μg of protein was injected onto the column. Thecolumn temperature was maintained at 40 °C, and the flow rate was set to 0.300 μL min^−1^ for the duration of the run. Solvent A (0.1% FA) and Solvent B (0.1% FA in ACN) were used as the chromatography solvents. The samples were run through the UHPLC System as follows: peptides were allowed to accumulate onto the trap column (Acclaim PepMap 100, C18, 75 μm x 2 cm, 3 μm, 100 Å column) for 10 min (during which the column was held at 2% Solvent B). The peptides were resolved by switching the trap column to be in-line with the separating column, quickly increasing the gradient to 5% B over 5 min and then applying a 95 min linear gradient from 5% B to 25% B. Subsequently, the gradient was increased from 35% B to 40% B over 25 min and then increased again from 40% B to 90% B over 5 min. The column was then cleaned with a sawtooth gradient to purge residual peptides between runs in a sequence.

A Thermo Q-Exactive HF-X Orbitrap mass spectrometer was used to analyze protein digests. A full MS scan in positive ion mode was followed by 20 data-dependent MS scans. The full MS scan was collected using a resolution of 120000 (@ m/z 200), an AGC target of 3E6, a maximum injection time of 64 ms, and a scan range from 350 to 1500 m/z. The data-dependent scans were collected with a resolution of 15000 (@ m/z 200), an AGC target of 1E5, a minimum AGC target of 8E3, a maximum injection time of 55 ms, and an isolation window of 1.4 m/z units. To dissociate precursors prior to their reanalysis by MS2, peptides were subjected to an HCD of 28% normalized collision energies. Fragments with charges of 1, 6, 7, or higher and unassigned were excluded from analysis, and a dynamic exclusion window of 30.0 s was used for the data-dependent scans. Mass tags were enabled with Δm of 2.00671 Th, 3.01007 Th, 4.01342 Th, and 6.02013 Th.

#### LC-MS/MS Data Analysis

Proteome Discoverer (PD) Software Suite (v2.4, Thermo Fisher) and the Minora Algorithm were used to analyze mass spectra and perform Label Free Quantification (LFQ) of detected peptides. Default settings for all analysis nodes were used except where specified. The data were searched against Escherichia coli (UP000000625, Uniprot) reference proteome database. For peptide identification, the PD MSFragger node was used, using a semi-tryptic search allowing up to 2 missed cleavages^72^. A precursor mass tolerance of 10 ppm was used for the MS1 level, and a fragment ion tolerance was set to 0.02 Da at the MS2 level. Peptide lengths between 7 and 50 amino acid residues was allowed with a peptide mass between 500 and 5000 Da. Additionally, a maximum charge state for theoretical fragments was set at 2. Oxidation of methionine and acetylation of the N-terminus were allowed as dynamic modifications while carbamidomethylation on cysteines was set as a static modification. Heavy isotope labeling (^13^C_6_) of Arginine and Lysine were allowed as dynamic modifications. The Philosopher PD node was used for FDR validation. Raw normalized extracted ion intensity data for the identified peptides were exported from the .pdResult file using a three-level hierarchy (protein > peptide group > consensus feature). These data were further processed utilizing custom Python analyzer scripts (available on GitHub, and described in depth previously^49,71^). Briefly, normalized ion counts were collected across the refolded replicates and the native replicates for each successfully identified peptide group. Effect sizes are the ratio of averages (reported in log2) and P-values (reported as −log10) were assessed using *t* tests with Welch’s correction for unequal population variances. Missing data are treated in a special manner. If a feature is not detected in all three native (or refolded) injections and is detected in all three refolded (or native) injections, we use those data, and fill the missing values with 1000 (the ion limit of detection for this mass analyzer); this peptide becomes classified as an all-or-nothing peptide. If a feature is not detected in one out of six injections, the missing value is dropped. Any other permutation of missing data (e.g., missing in two injections) results in the quantification getting discarded. In many situations, our data provide multiple independent sets of quantifications for the same peptide group. This happens most frequently because the peptide is detected in multiple charge states or as a heavy isotopomer. In this case, we calculate effect size and P-value for all features that map to the same peptide group. If the features all agree with each other in sign, they are combined: the quantification associated with the median amongst available features is used and the P-values are combined with Fisher’s method. If the features disagree with each other in sign, the P-value is set to 1. Coefficients of variation (CV) for the peptide abundance in the three replicate refolded samples are also calculated. Analyzer returns a file listing all the peptides that can be confidently quantified, and provides their effect-size, P-value, refolded CV, proteinase K site (if half-tryptic), and associated protein metadata.

**Figure S1.**
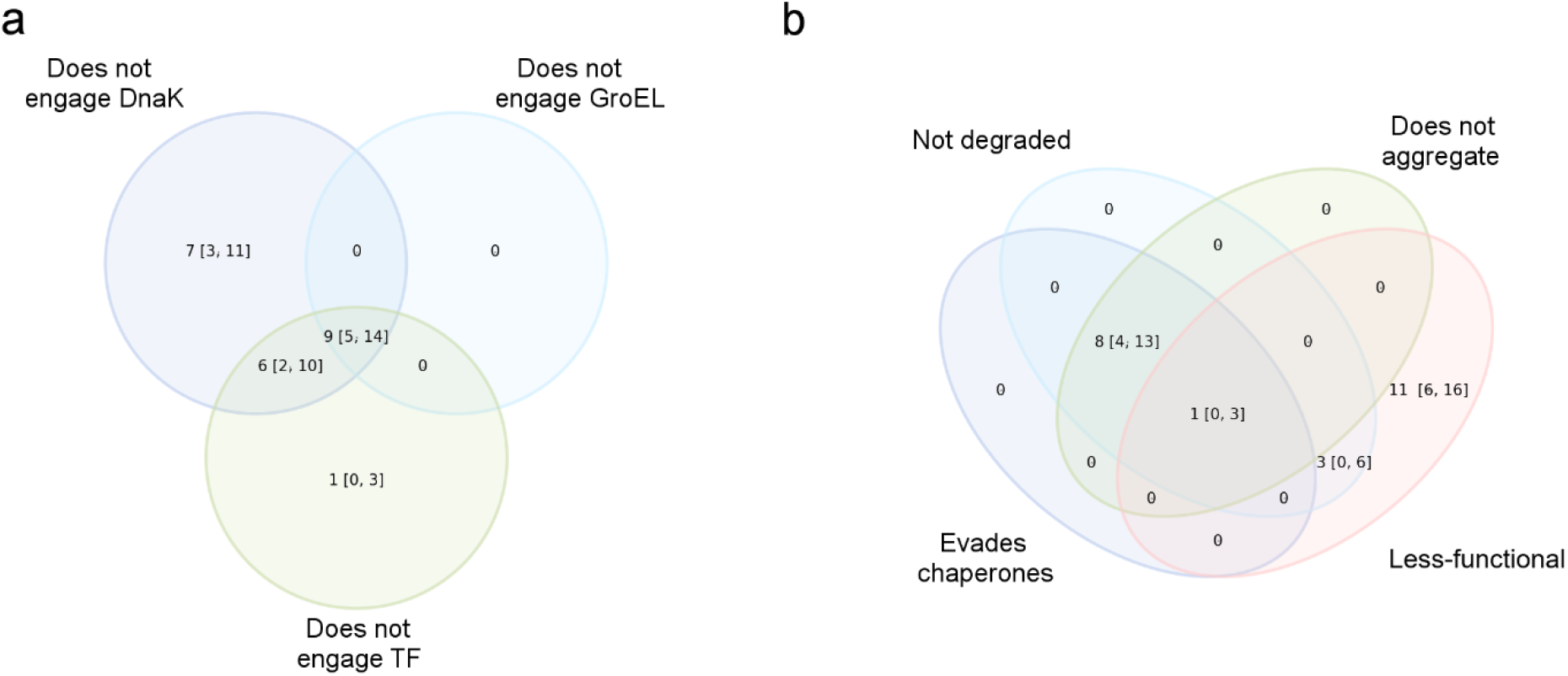
Venn diagrams showing the overlap between Luciferase trajectories that are misfolded and (a) predicted to evade DnaK, GroEL, and TF or (b) predicted not to aggregate, not to be degraded, to evade all chaperones, and to remain non-functional. Error bars are 95% confidence intervals from bootstrapping 10^6^ times.

**Figure S2.**
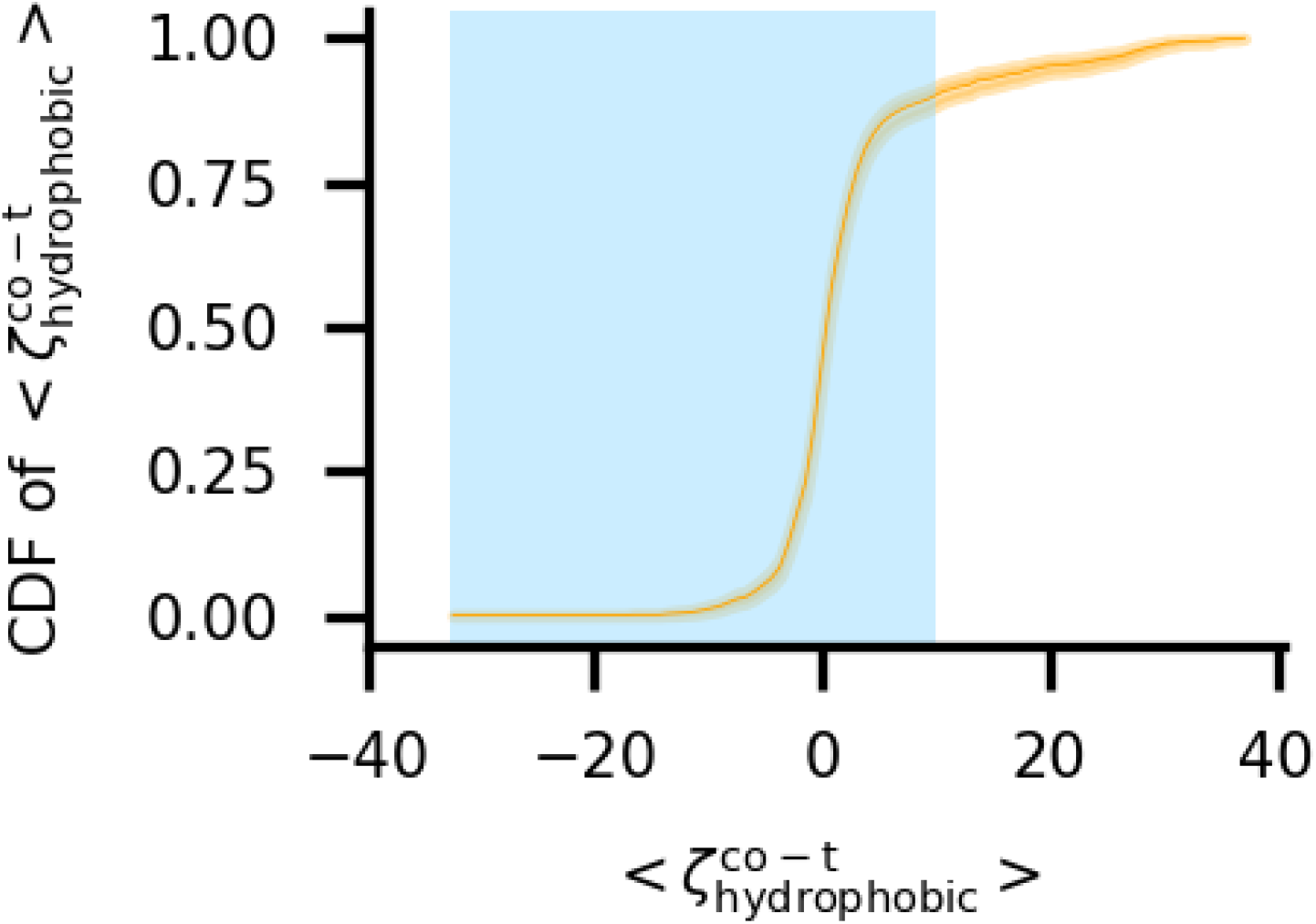
Cumulative distribution function of 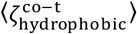 over the subset of 1,053 misfolded trajectories for which it was computed (see Methods). The blue shaded region indicates the subset of values taken to indicate no significant increase in trigger factor interactions relative to the folded population. The height of the CDF indicates the 95% confidence intervals.

**Figure S3.**
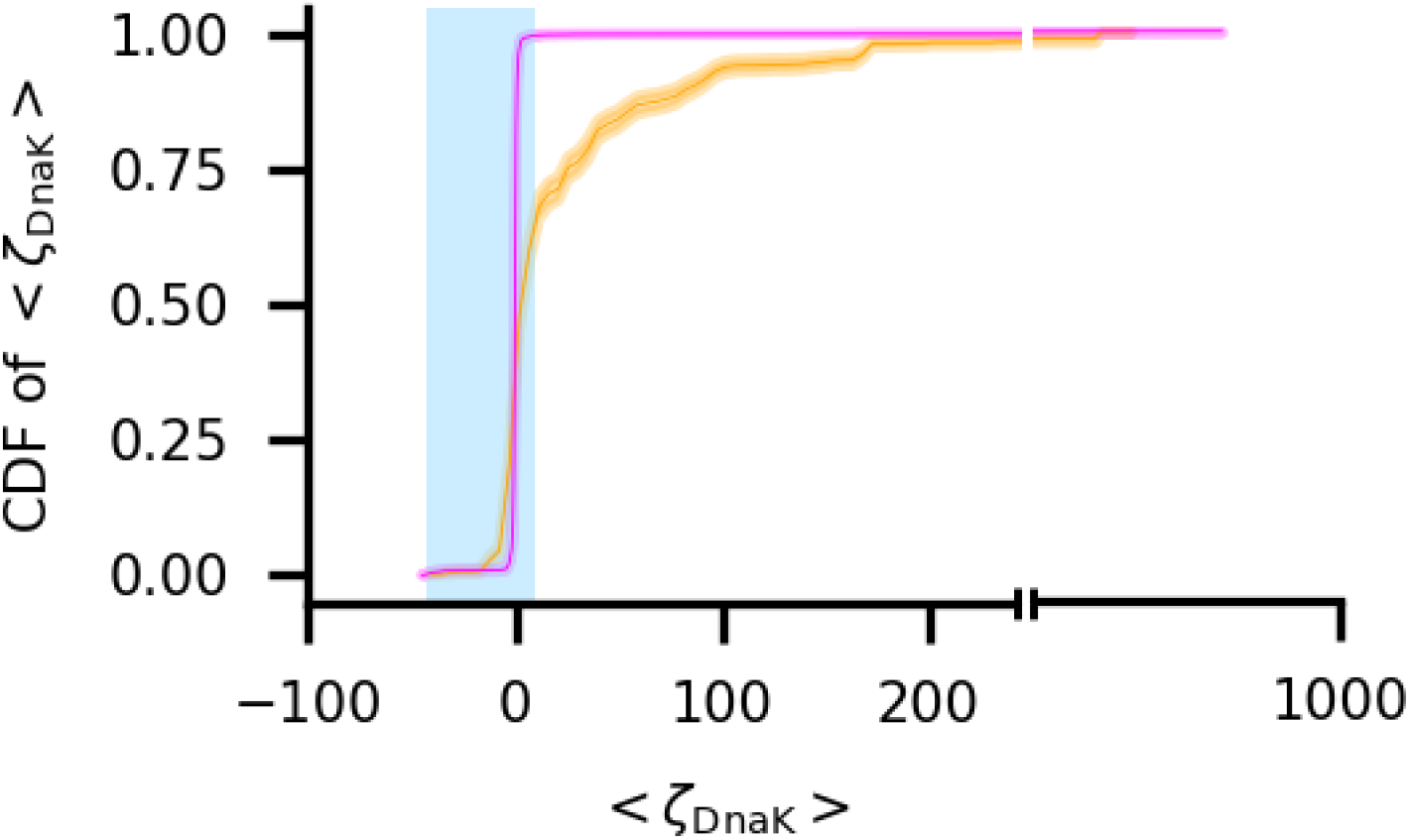
Cumulative distribution function of 〈*ζ*_DnaK_〉 over the subset of 1,631 misfolded (orange) and 4,469 folded (magenta) trajectories. The blue shaded region indicates the subset of values taken to indicate no significant increase in DnaK interactions relative to the native-state population. The height of the CDFs indicates the 95% confidence intervals.

**Figure S4.**
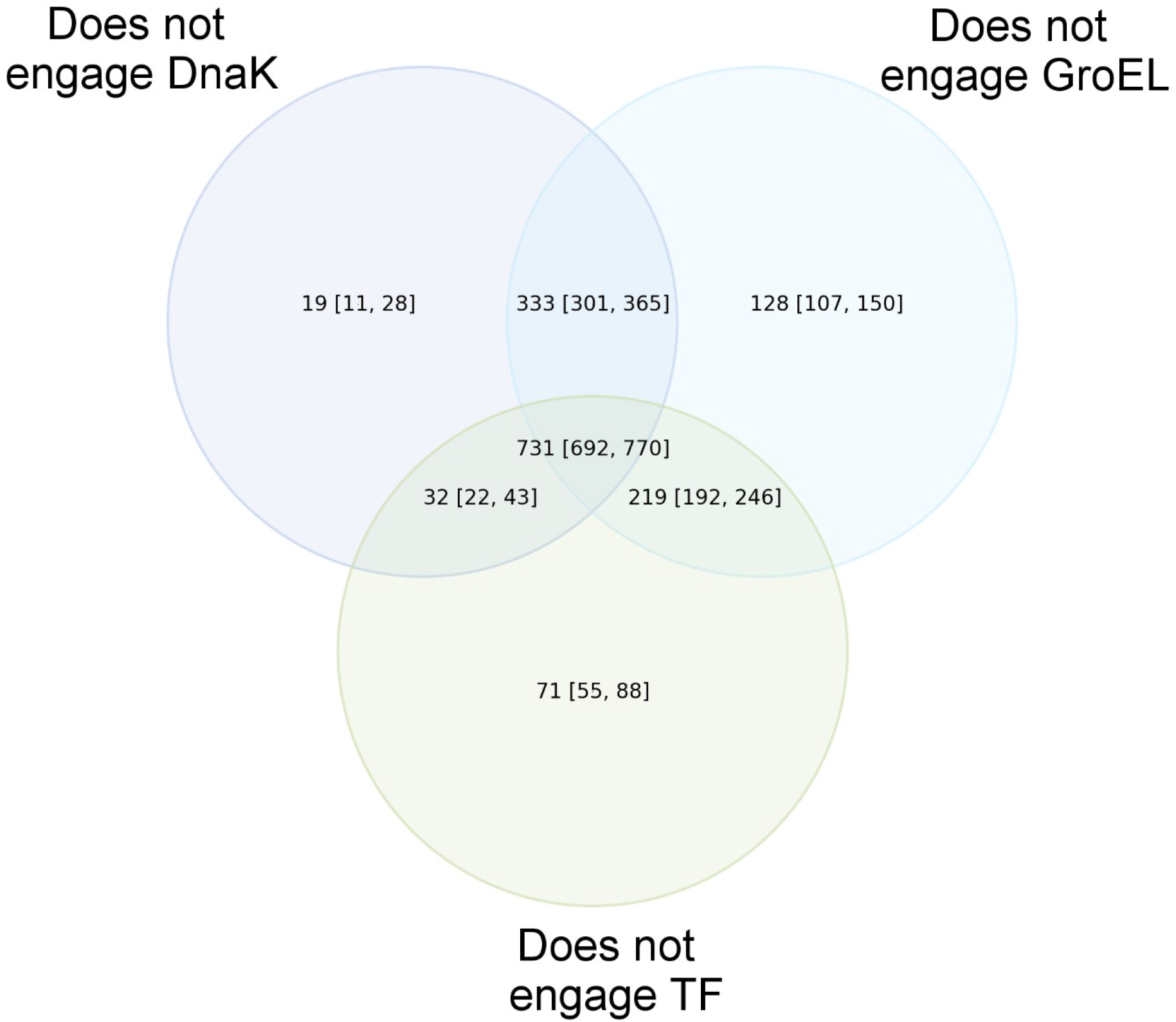
Venn diagram showing the overlap between those trajectories that are predicted not to engage TF, DnaK, or GroEL/GroES. A total of 1,533 trajectories evade at least on chaperone. Error bars are 95% confidence intervals from bootstrapping 10^6^ times.

**Figure S5.**
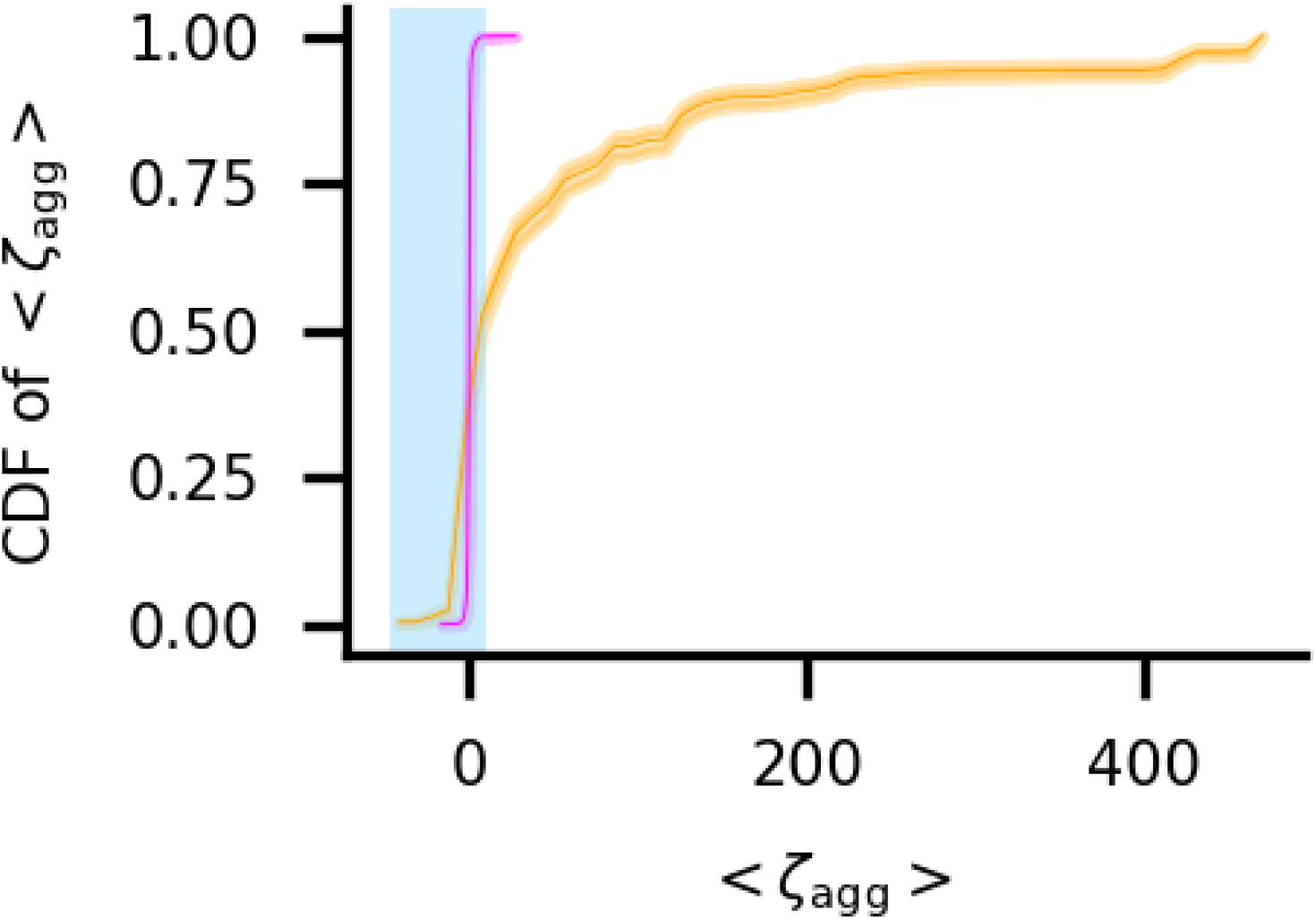
Cumulative distribution function of 〈*ζ*_Agg_〉 over the subset of 1,631 misfolded (orange) and 4,469 folded (magenta) trajectories. The blue shaded region indicates the subset of values taken to indicate no significant increase in aggregation propensity relative to the native state simulations. The height of the CDFs indicates the 95% confidence intervals.

**Figure S6.**
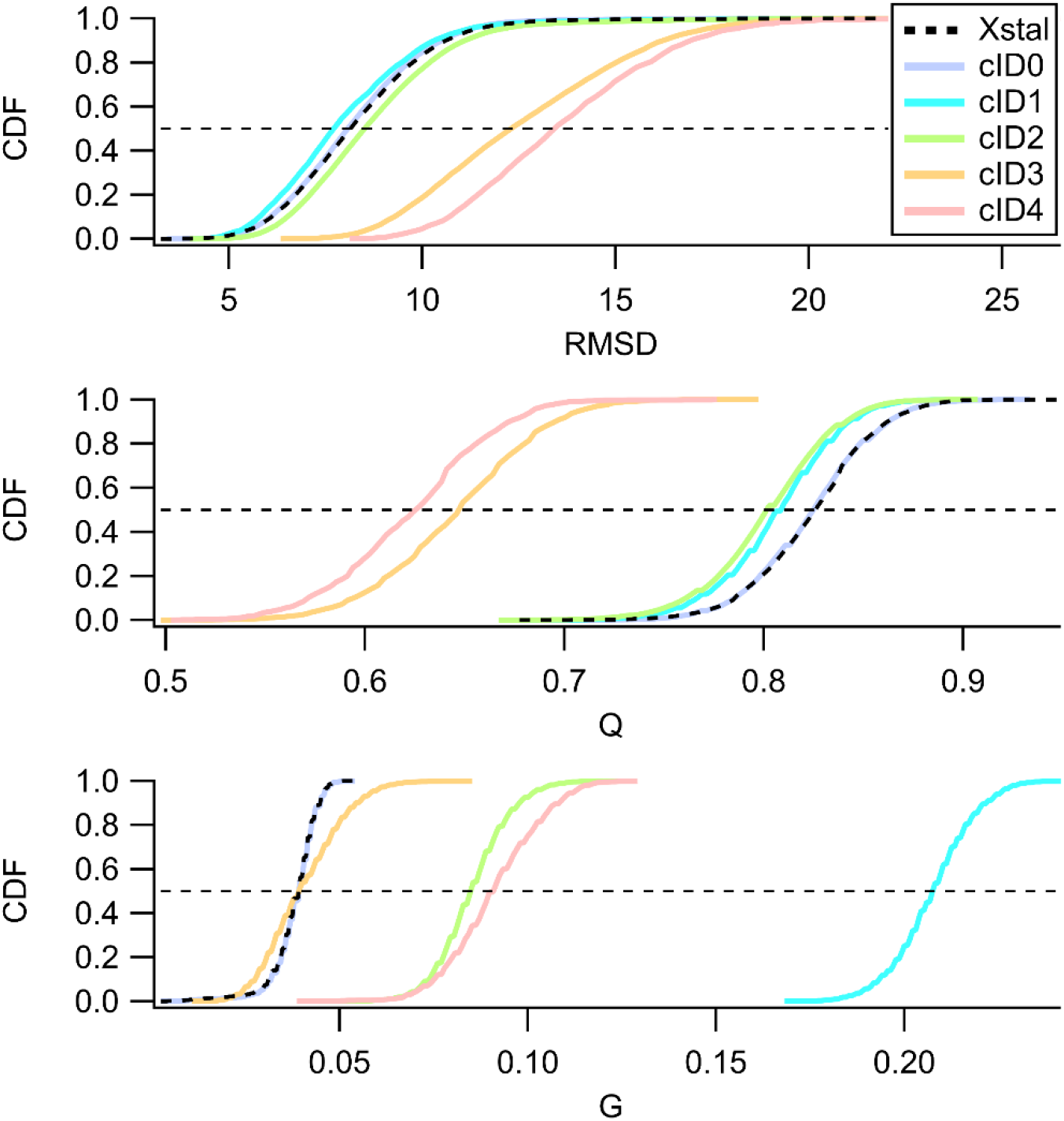
The utility of the change in entanglement metric (*G*) for detecting structural perturbations not apparent by fraction of native contacts (*Q*) or root mean square deviation (RMSD) is exemplified by examining clusters of the trajectories for aminoimidazole ribonucleotide synthetase (PDB: 1CLI) relative to the cluster of trajectories started from the reference state. The 50 trajectories obtained from synthesis simulations can be clustered into 5 separate clusters characterized by the 〈*Q*_mode_〉 and the corrected discrete distribution of a give type of change in entanglement (Eq. 7). cID0 contains 8/50 trajectories all of which terminate to the native state with no changes in entanglement. cID1 contains 3/50 trajectories that are near native like (i.e. 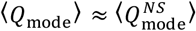) that show appreciable gain and loss of entanglements as well as pure switch in chirality. cID2 contains 32/50 trajectories that show appreciable gain and loss of entanglements with less near native like conformations (i.e. 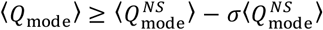). cID3 contains 6/50 trajectories that show non-native like conformations (i.e. 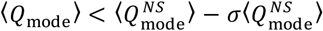) with appreciable gain and loss of entanglements. cID4 contains 1/50 trajectories that show non-native like conformations with appreciable gain and loss of entanglements and pure change in chirality. For the clusters which contain near native like conformations and changes in entanglement of any kind (cID1 light blue and cID2) *Q* and RMSD fail to distinguish different clusters while G does not. For those clusters that contain non-native like conformations and changes in entanglement of any kind (cID3 & cID4) *Q* and RMSD are much more reliable.

**Table S1.** Coarse-grain model parameters for PDB ID 4G36

| PDB ID | Protein name | Domain or interface | Structural class | $\eta$ used in model building |
| --- | --- | --- | --- | --- |
| 4G36 | Firefly Luciferase | Domain 1:1-12; 356-437 | $\beta$ | 1.442 |
| | | Domain 2:13-52; 212-355 | $\alpha/\beta$ | 1.114 |
| | | Domain 3: 53-211 | $\alpha/\beta$ | 1.114 |
| | | Domain 4: 438-550 | $\alpha/\beta$ | 1.916 |
|  |  | Interface 1\|2 | - | 1.235 |
|  |  | Interface 1\|3 | - | 1.235 |
|  |  | Interface 1\|4 | - | 1.507* |
|  |  | Interface 2\|3 | - | 1.235 |
|  |  | Interface 2\|4 | - | 1.235 |
|  |  | Interface 3\|4 | - | 1.235 |

\*indicates interface is unstable at all tested values of $\eta$ and so median interface value from set of stable interfaces is used for all simulations

**Table S2.**
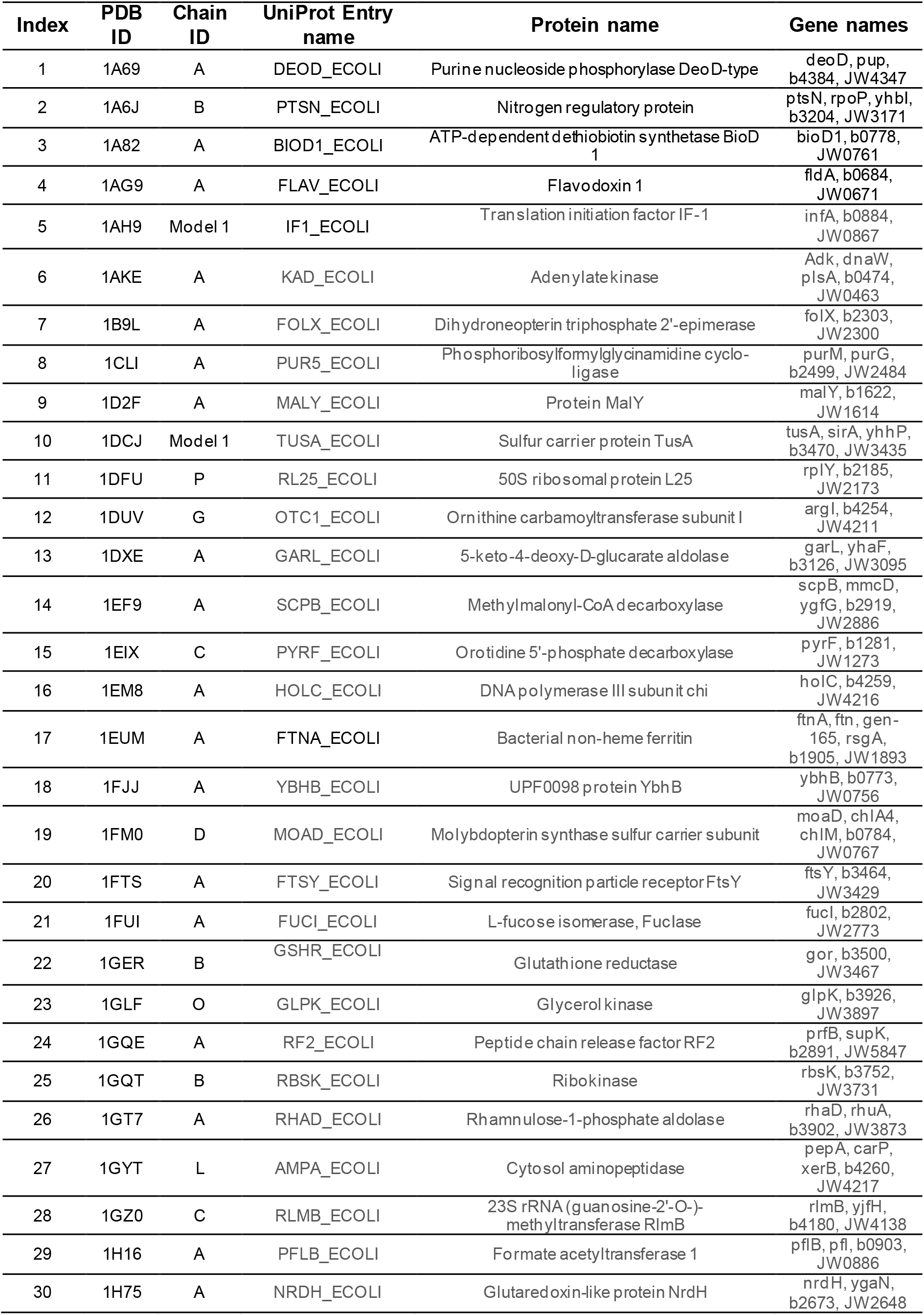
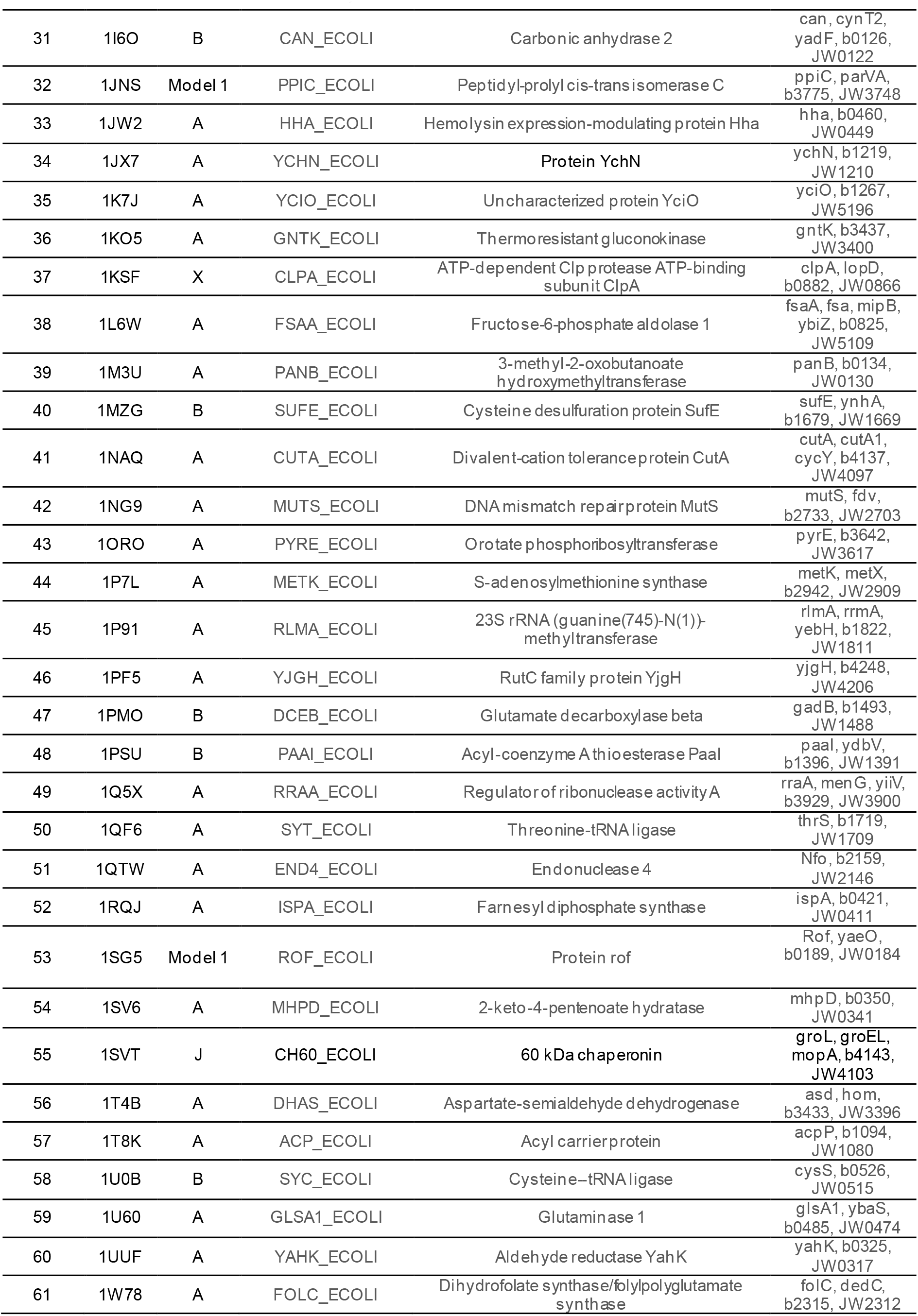
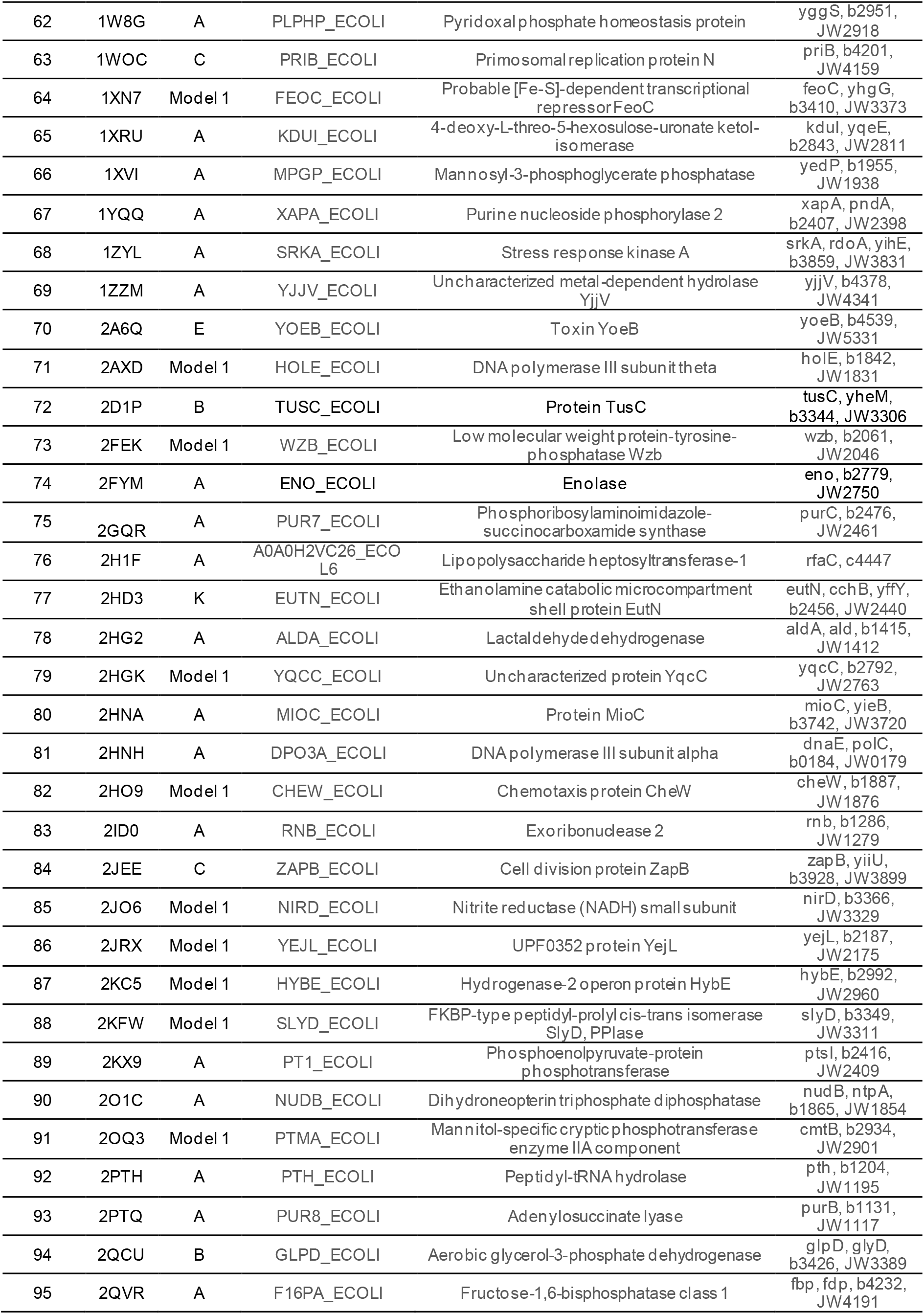
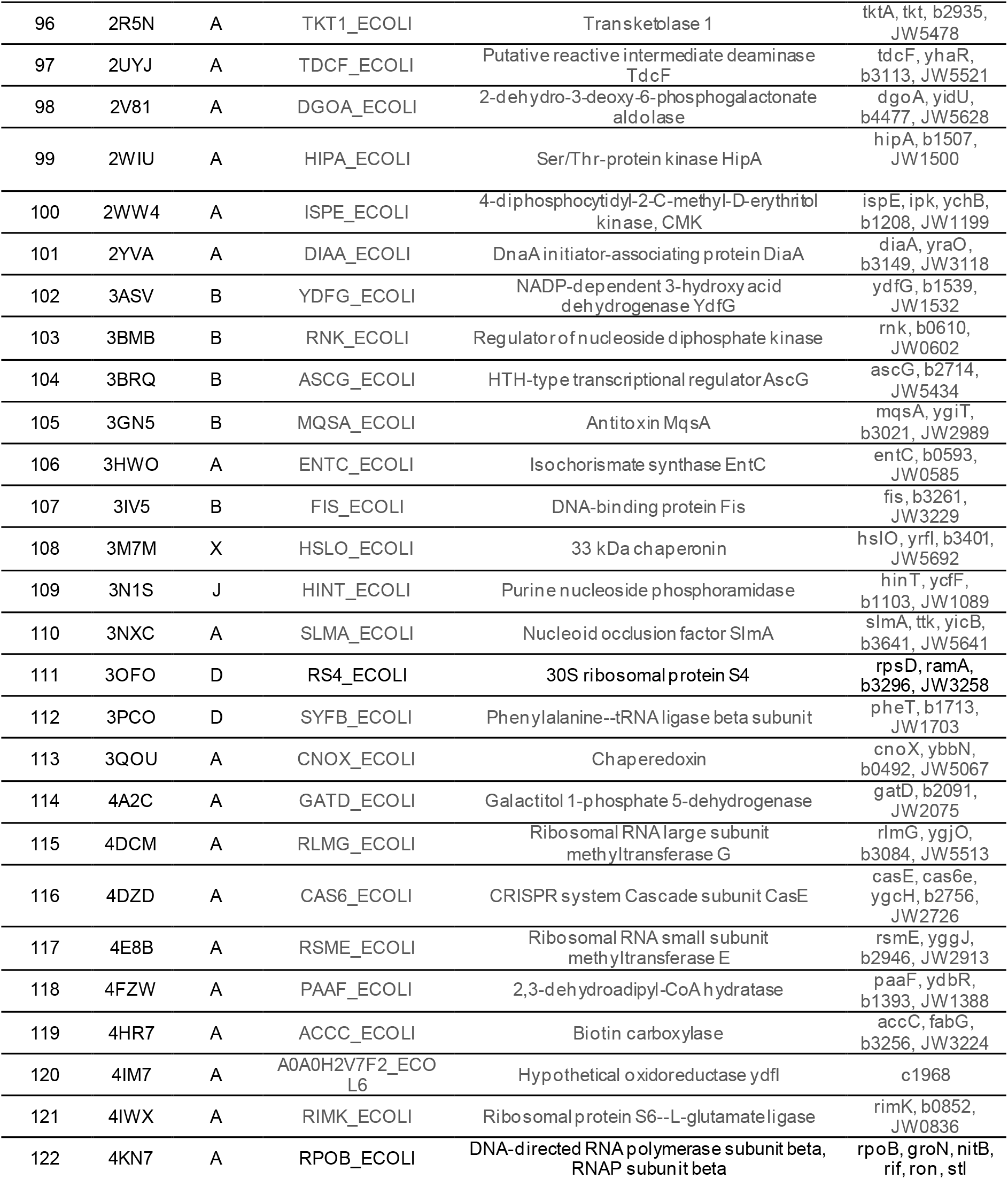
*E. coli* proteome dataset information

**Table S3.**
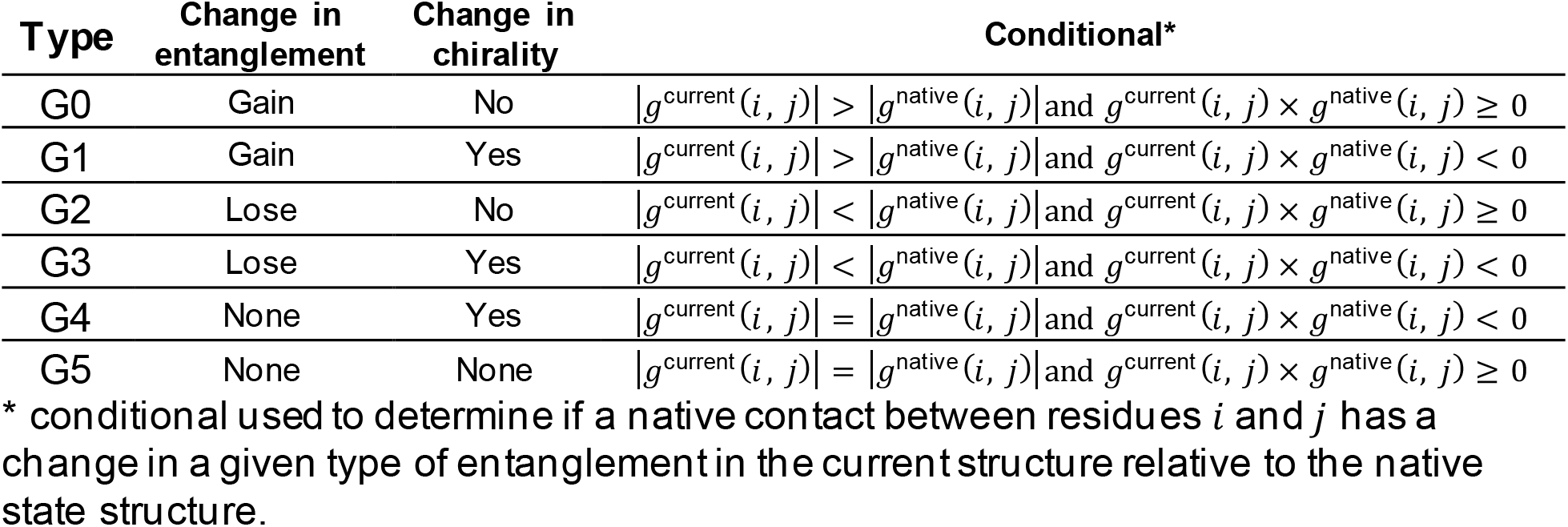
Definitions of entanglement types

**Table S4.**
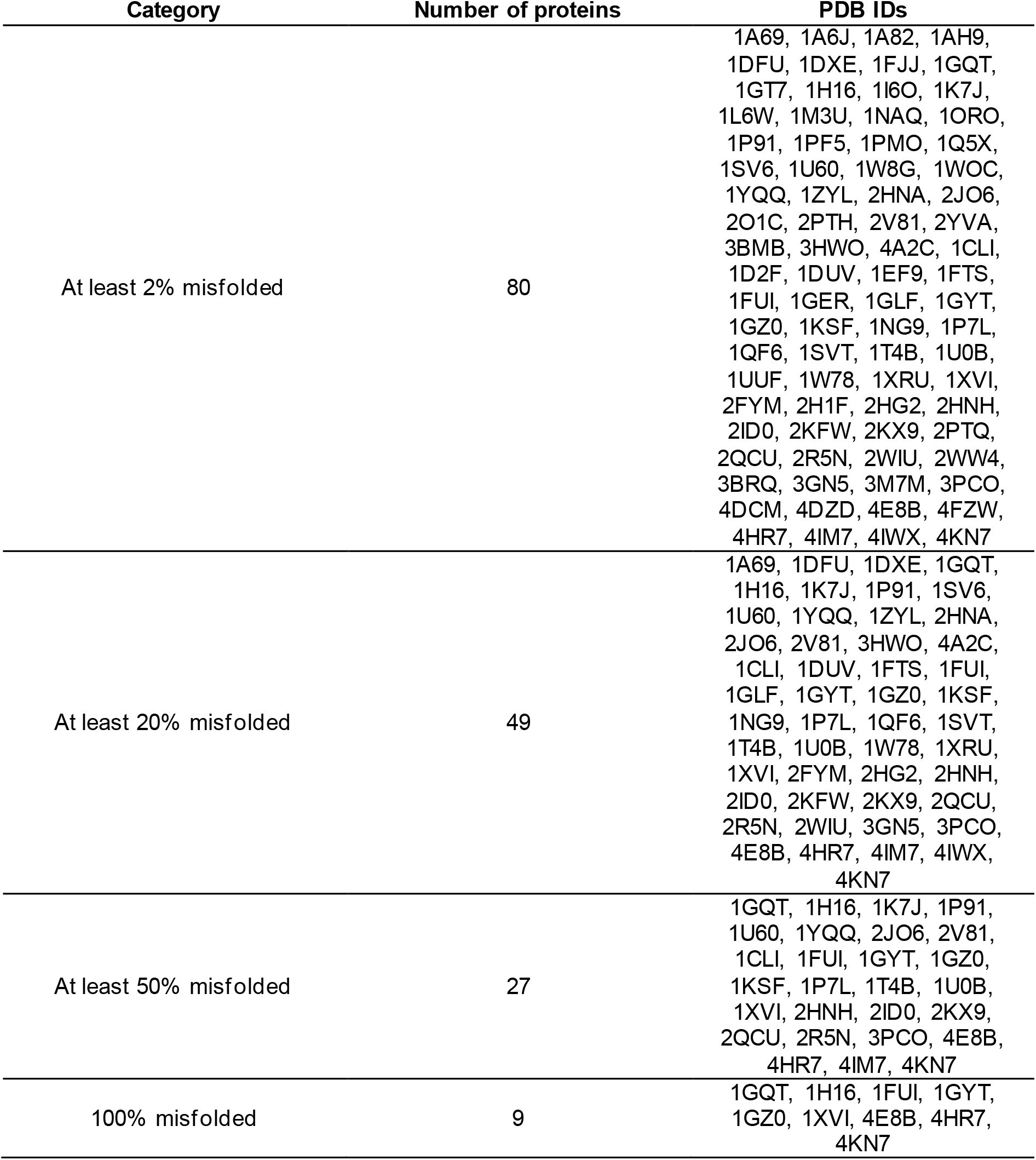
Proteins categorized by percent of trajectories misfolded based

**Table S5.**
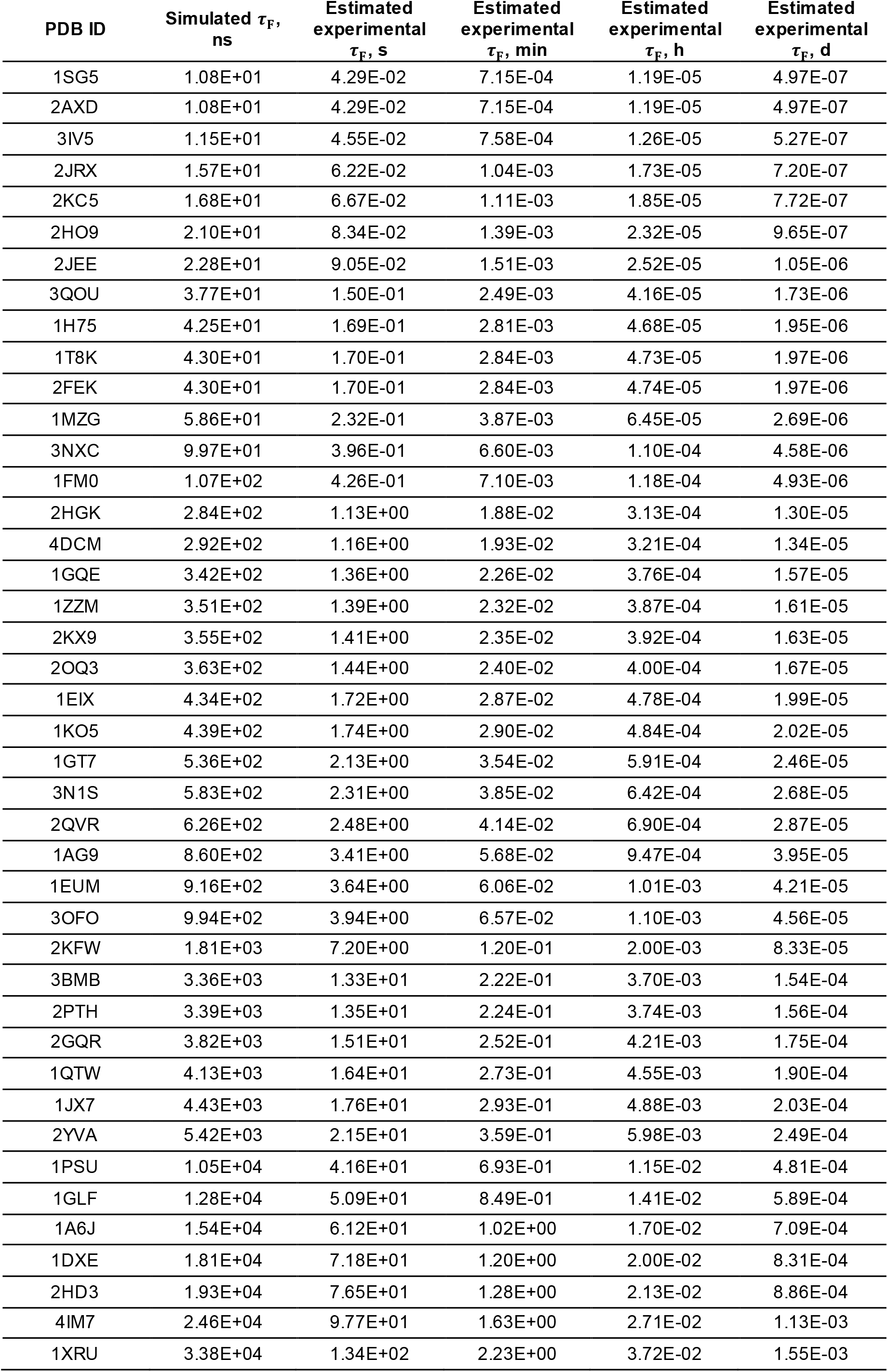
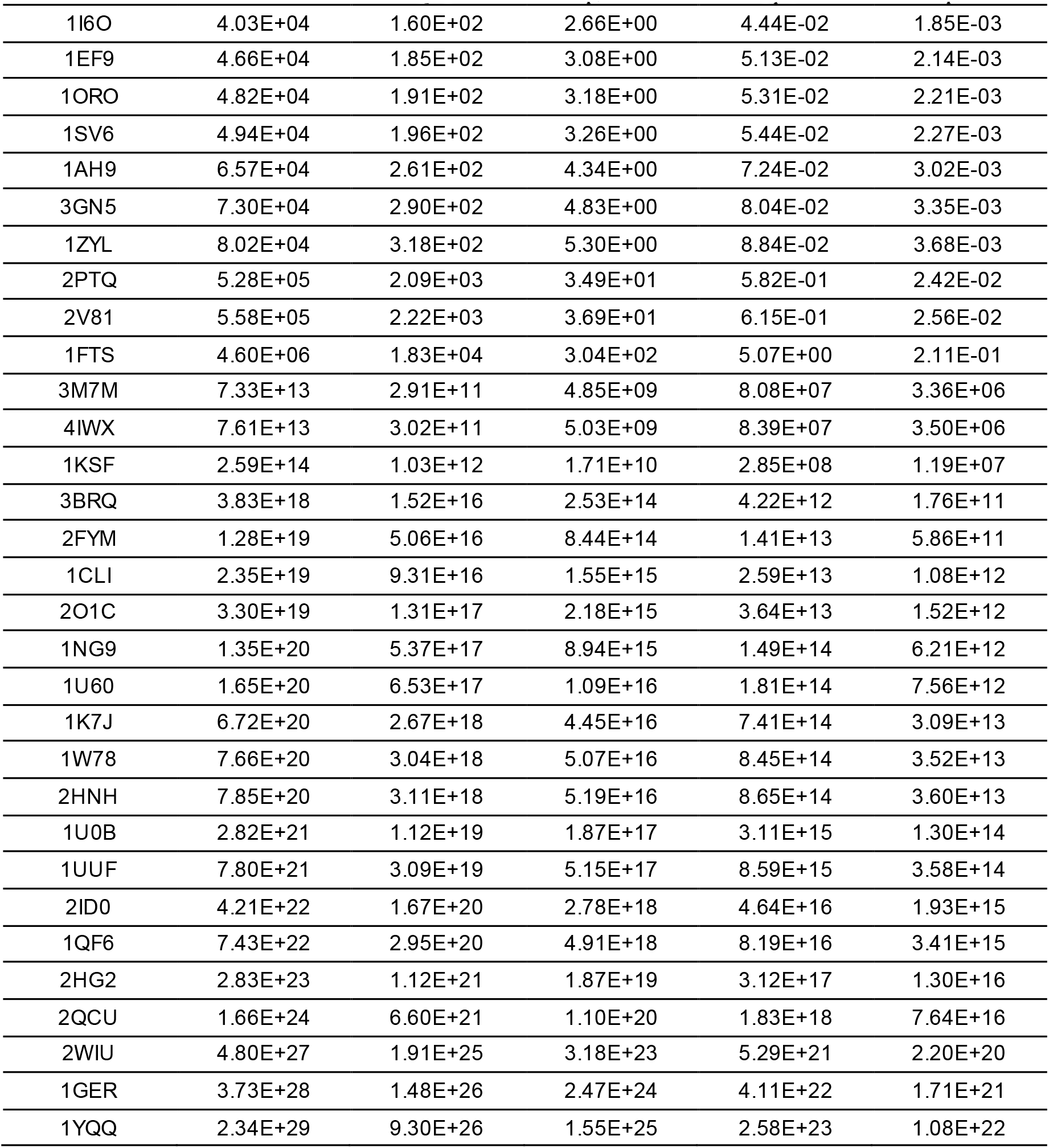
Extrapolated folding times for the 73 proteins with a reliable estimate

**Table S6.**
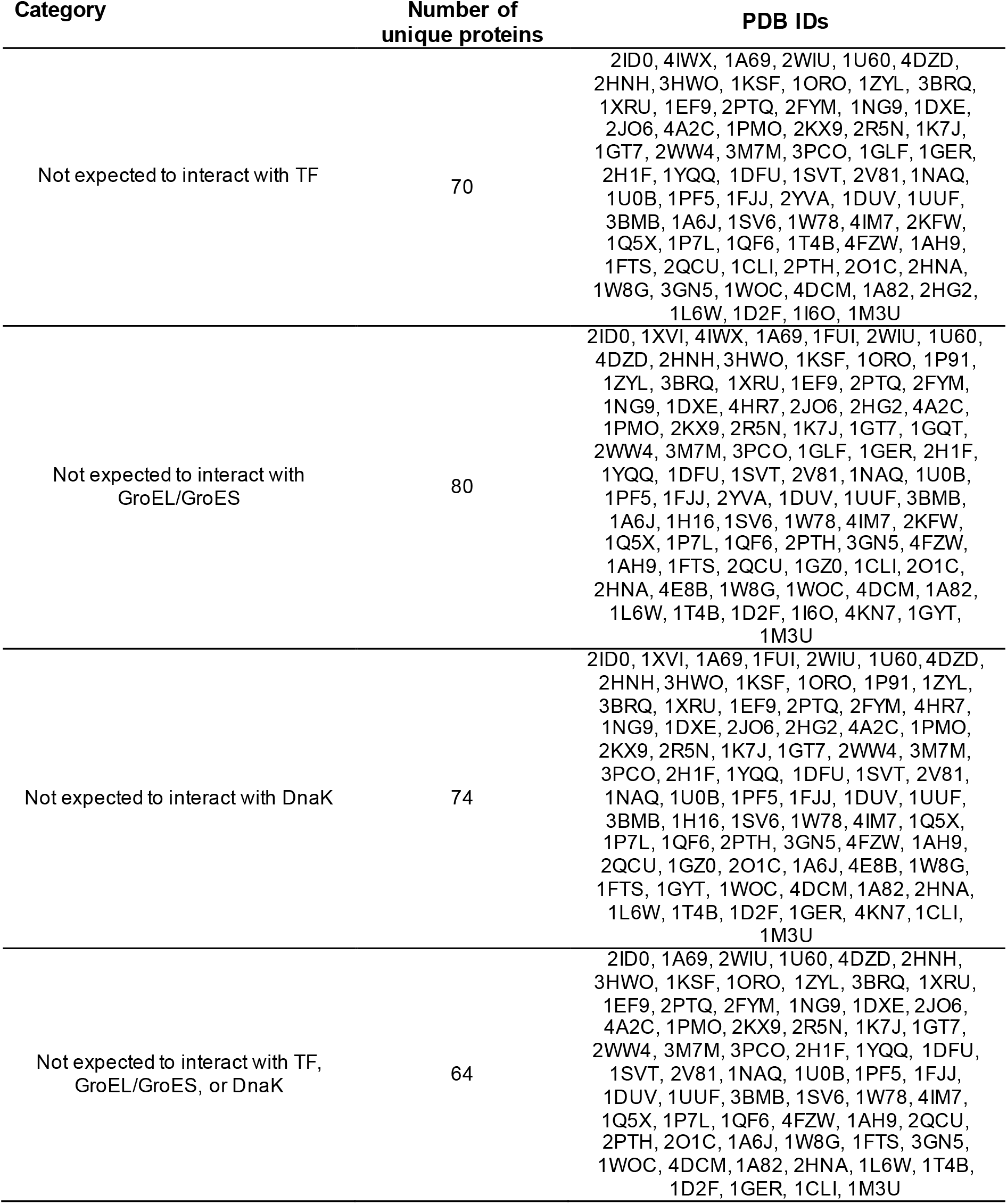
Proteins with at least one misfolded trajectory expected to bypass chaperones

**Table S7.**
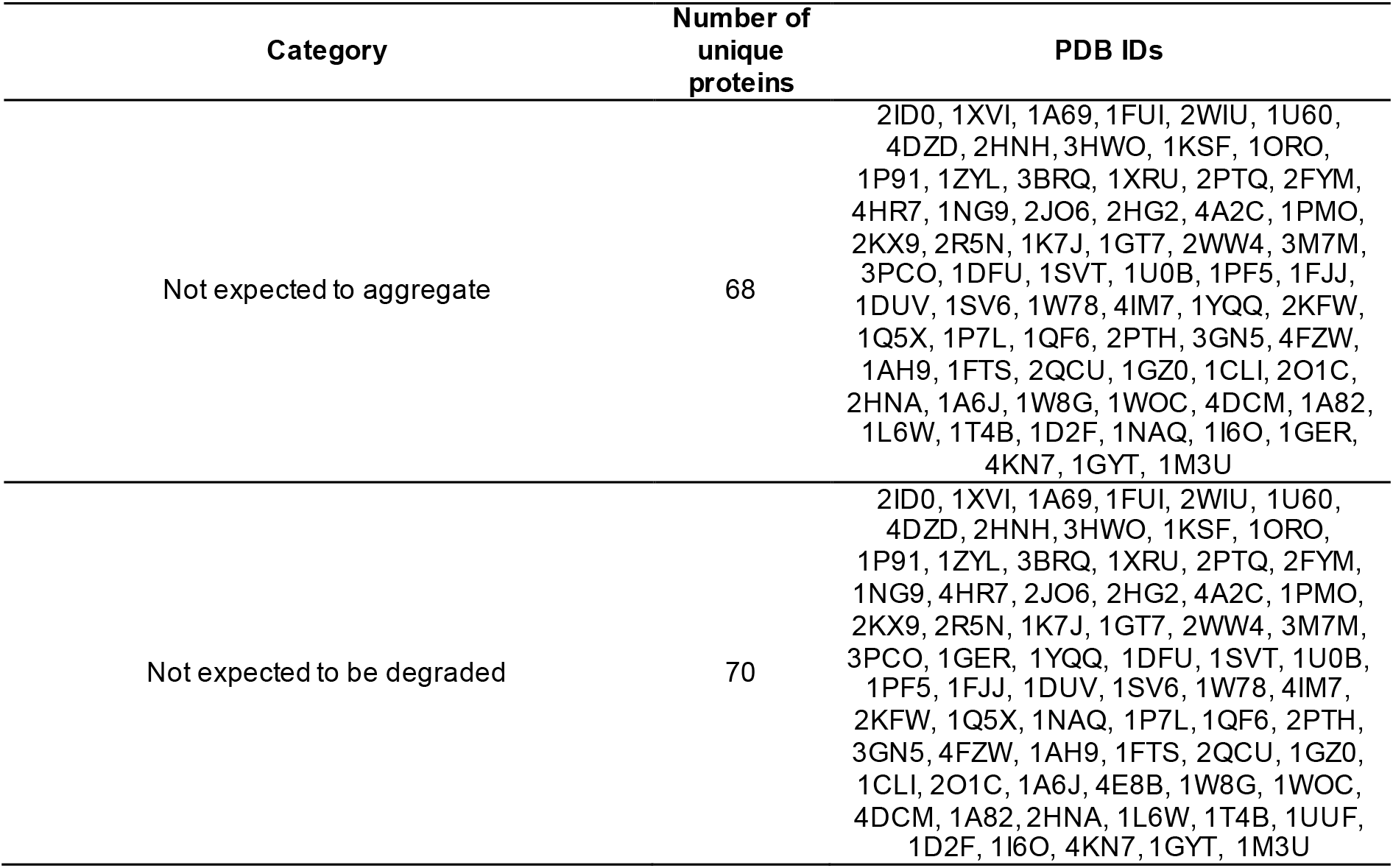
Proteins with at least one misfolded trajectory expected not to aggregate or be degraded

**Table S8.**
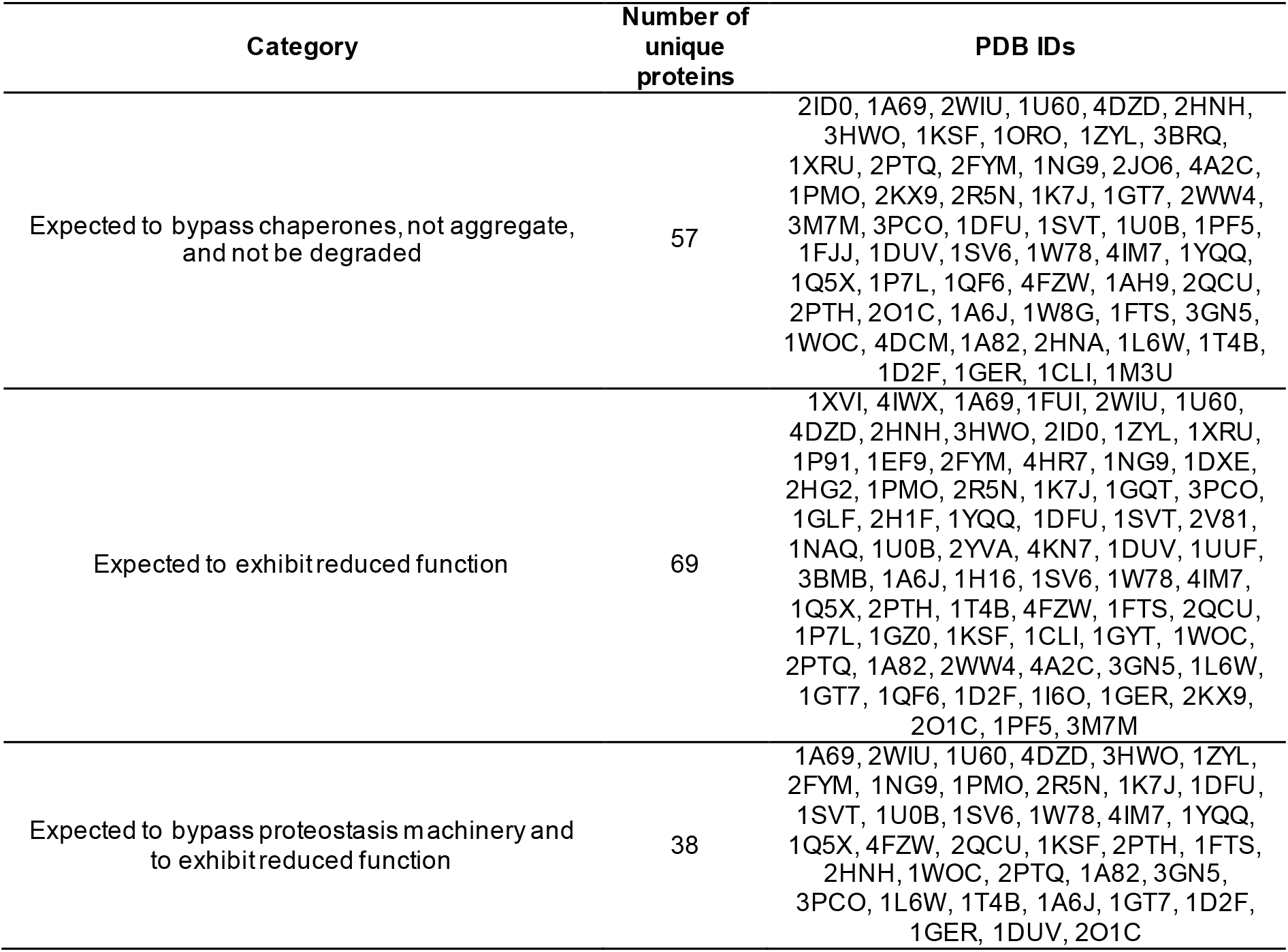
Proteins with misfolded conformations expected to bypass proteostasis machinery and remain soluble but less functional

**Table S9.** Percent of native state simulation frames with metric ≥10% averaged over set of 122 *E. coli* proteins

| **Metric** | **Mean [95% CIs]\*, %** | **Range (low-high), %** |
| --- | --- | --- |
| $\zeta_{\text{hydrophobic}}(t)$, | 6.32 [5.44, 7.27] | 0.08 – 32.31 |
| $\zeta_{\text{DnaK}}(t)$ | 14.74 [13.03, 16.47] | 0.17 – 38.35 |
| $\zeta_{\text{agg}}(t)$ | 11.74 [10.32, 13.22] | 0.60 – 39.75 |
| $\chi_{\text{func}}$ | 24.58 [22.97, 26.28] | 9.52 – 55.47 |

\*confidence intervals calculated from bootstrapping $10^6$ times

**Table S10.**
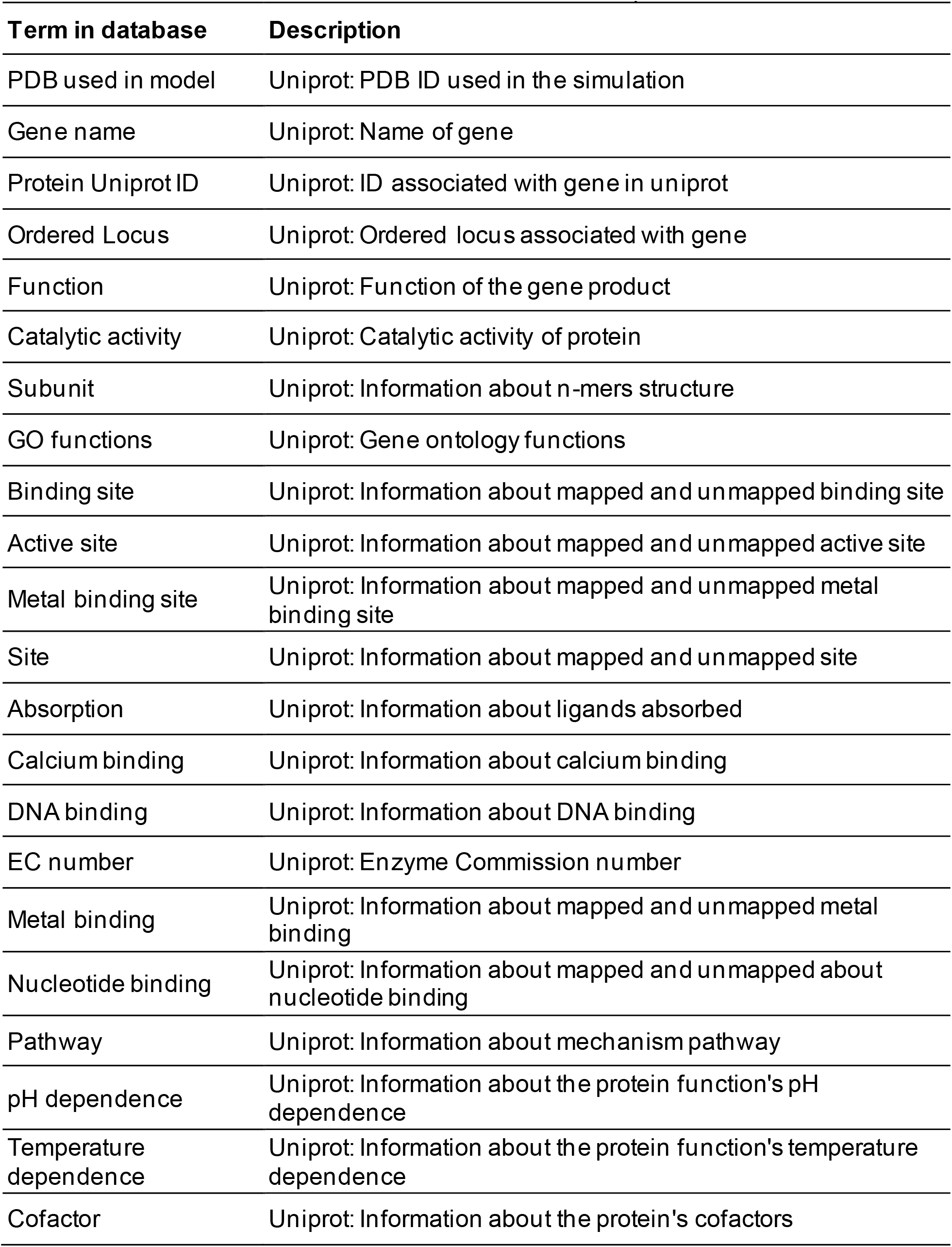
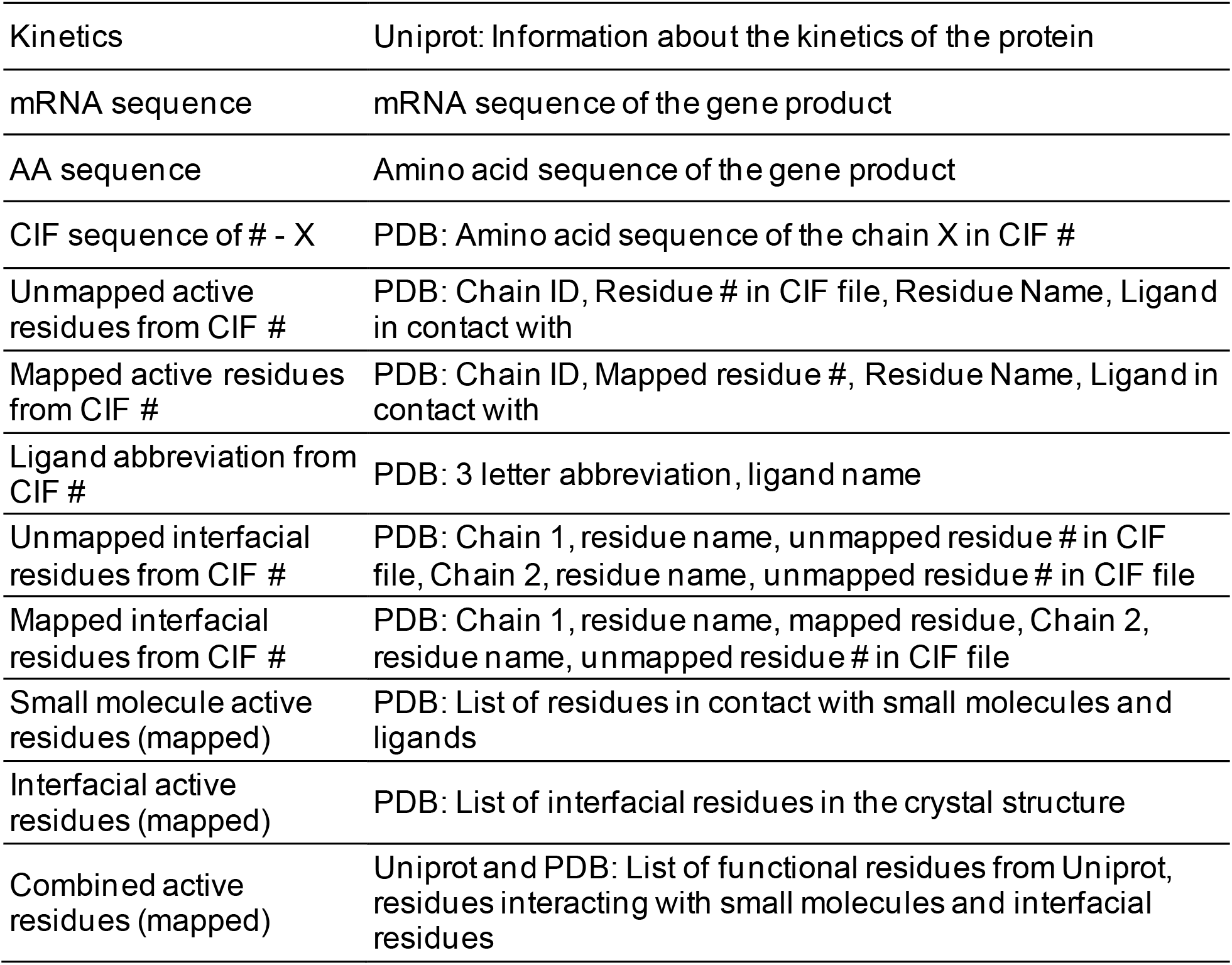
Functional residues database terms and descriptions

| Term in database | Description |
| --- | --- |
| PDB used in model | Uniprot: PDB ID used in the simulation |
| Gene name | Uniprot: Name of gene |
| Protein Uniprot ID | Uniprot: ID associated with gene in uniprot |
| Ordered Locus | Uniprot: Ordered locus associated with gene |
| Function | Uniprot: Function of the gene product |
| Catalytic activity | Uniprot: Catalytic activity of protein |
| Subunit | Uniprot: Information about n-mers structure |
| GO functions | Uniprot: Gene ontology functions |
| Binding site | Uniprot: Information about mapped and unmapped binding site |
| Active site | Uniprot: Information about mapped and unmapped active site |
| Metal binding site | Uniprot: Information about mapped and unmapped metal binding site |
| Site | Uniprot: Information about mapped and unmapped site |
| Absorption | Uniprot: Information about ligands absorbed |
| Calcium binding | Uniprot: Information about calcium binding |
| DNA binding | Uniprot: Information about DNA binding |
| EC number | Uniprot: Enzyme Commission number |
| Metal binding | Uniprot: Information about mapped and unmapped metal binding |
| Nucleotide binding | Uniprot: Information about mapped and unmapped about nucleotide binding |
| Pathway | Uniprot: Information about mechanism pathway |
| pH dependence | Uniprot: Information about the protein function's pH dependence |
| Temperature dependence | Uniprot: Information about the protein function's temperature dependence |
| Cofactor | Uniprot: Information about the protein's cofactors |

**Table S10** continued
| Term in database | Description |
| --- | --- |
| Kinetics | Uniprot: Information about the kinetics of the protein |
| mRNA sequence | mRNA sequence of the gene product |
| AA sequence | Amino acid sequence of the gene product |
| CIF sequence of # - X | PDB: Amino acid sequence of the chain X in CIF # |
| Unmapped active residues from CIF # | PDB: Chain ID, Residue # in CIF file, Residue Name, Ligand in contact with |
| Mapped active residues from CIF # | PDB: Chain ID, Mapped residue #, Residue Name, Ligand in contact with |
| Ligand abbreviation from CIF # | PDB: 3 letter abbreviation, ligand name |
| Unmapped interfacial residues from CIF # | PDB: Chain 1, residue name, unmapped residue # in CIF file, Chain 2, residue name, unmapped residue # in CIF file |
| Mapped interfacial residues from CIF # | PDB: Chain 1, residue name, mapped residue, Chain 2, residue name, unmapped residue # in CIF file |
| Small molecule active residues (mapped) | PDB: List of residues in contact with small molecules and ligands |
| Interfacial active residues (mapped) | PDB: List of interfacial residues in the crystal structure |
| Combined active residues (mapped) | Uniprot and PDB: List of functional residues from Uniprot, residues interacting with small molecules and interfacial residues |

**Table S11.**
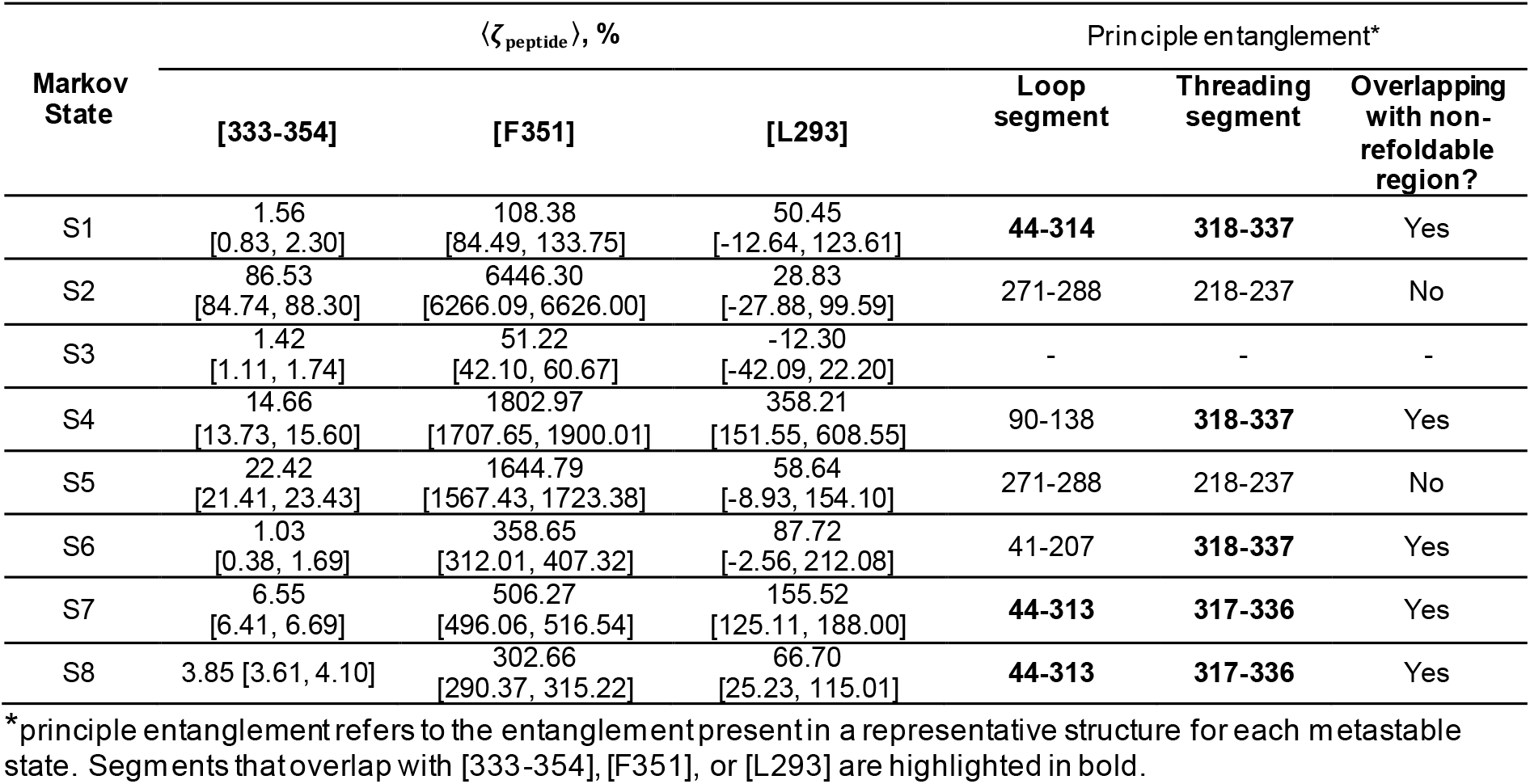
Values of 〈*ζ*_peptide_〉 and locations of principle entanglements by metastable state

